# Structural mimicry confers robustness in the cyanobacterial circadian clock

**DOI:** 10.1101/2020.06.17.158394

**Authors:** Joel Heisler, Jeffrey A. Swan, Joseph G. Palacios, Cigdem Sancar, Dustin C. Ernst, Rebecca K. Spangler, Clive R. Bagshaw, Sarvind Tripathi, Priya Crosby, Susan S. Golden, Carrie L. Partch, Andy LiWang

## Abstract

The histidine kinase SasA enhances robustness of circadian rhythms in the cyanobacterium *S. elongatus* by temporally controlling expression of the core clock components, *kaiB* and *kaiC*. Here we show that SasA also engages directly with KaiB and KaiC proteins to regulate the period and enhance robustness of the reconstituted circadian oscillator *in vitro*, particularly under limiting concentrations of KaiB. In contrast to its role regulating gene expression, oscillator function does not require SasA kinase activity; rather, SasA uses structural mimicry to cooperatively recruit the rare, fold-switched conformation of KaiB to the KaiC hexamer to form the nighttime repressive complex. Cooperativity gives way to competition with increasing concentrations of SasA to define a dynamic window by which SasA directly modulates clock robustness.

**One Sentence Summary:** SasA controls the assembly of clock protein complexes through a balance of cooperative and competitive interactions.

## Main Text

The core circadian clock genes *kaiA, kaiB, and kaiC*, are essential for rhythmic gene expression in cyanobacteria (*1*) and their proteins generate a ~24-hour rhythm of phosphorylation of KaiC *in vivo* (*2*) that persists in the absence of transcription or translation (*3*). Reconstitution of this post-translational oscillator (PTO) *in vitro* (*4*) has led to deep insight into the structural and kinetic mechanisms that regulate formation of the clock protein assemblies and underlie cyanobacterial circadian rhythms. KaiC autophosphorylation is stimulated by KaiA during the day (*5–7*), whereas autodephosphorylation is favored at night when KaiB binds to KaiC and sequesters KaiA (*8, 9*) (**Fig. 1A**). Although these three proteins alone can establish a circadian oscillation *in vitro*, two histidine protein kinases contribute to the complex *in vivo:* SasA and CikA, which coordinate a vast program of circadian gene expression executed by the response regulator, RpaA (*10*). The N-terminal thioredoxin-like domain of SasA is structurally homologous to a rare fold-switched monomer state of KaiB, and directly competes with this form of KaiB for binding to KaiC (*11*, *12*). This structural similarity and binding competition raised the possibility that SasA helps to regulate formation of the nighttime repressive complex, which is known to be restricted by (*i*) the phosphorylation state of the KaiC CII domain (*13*), (*ii*) the intrinsically slow rate of ATP hydrolysis by the KaiC CI domain (*14, 15*), and (*iii*) the rare conversion of KaiB to the its fold-switched form that is competent to bind KaiC (*11*). Similarly, CikA and KaiA bind to the same site on the KaiBC complex (*9*), and this competition shortens the period (*11*) and compensates for diminished concentrations of KaiA in the PTO *in vitro* (*16*), suggesting that output kinases fortify robustness of the PTO by modulating Kai protein interactions directly.

**Figure 1:**
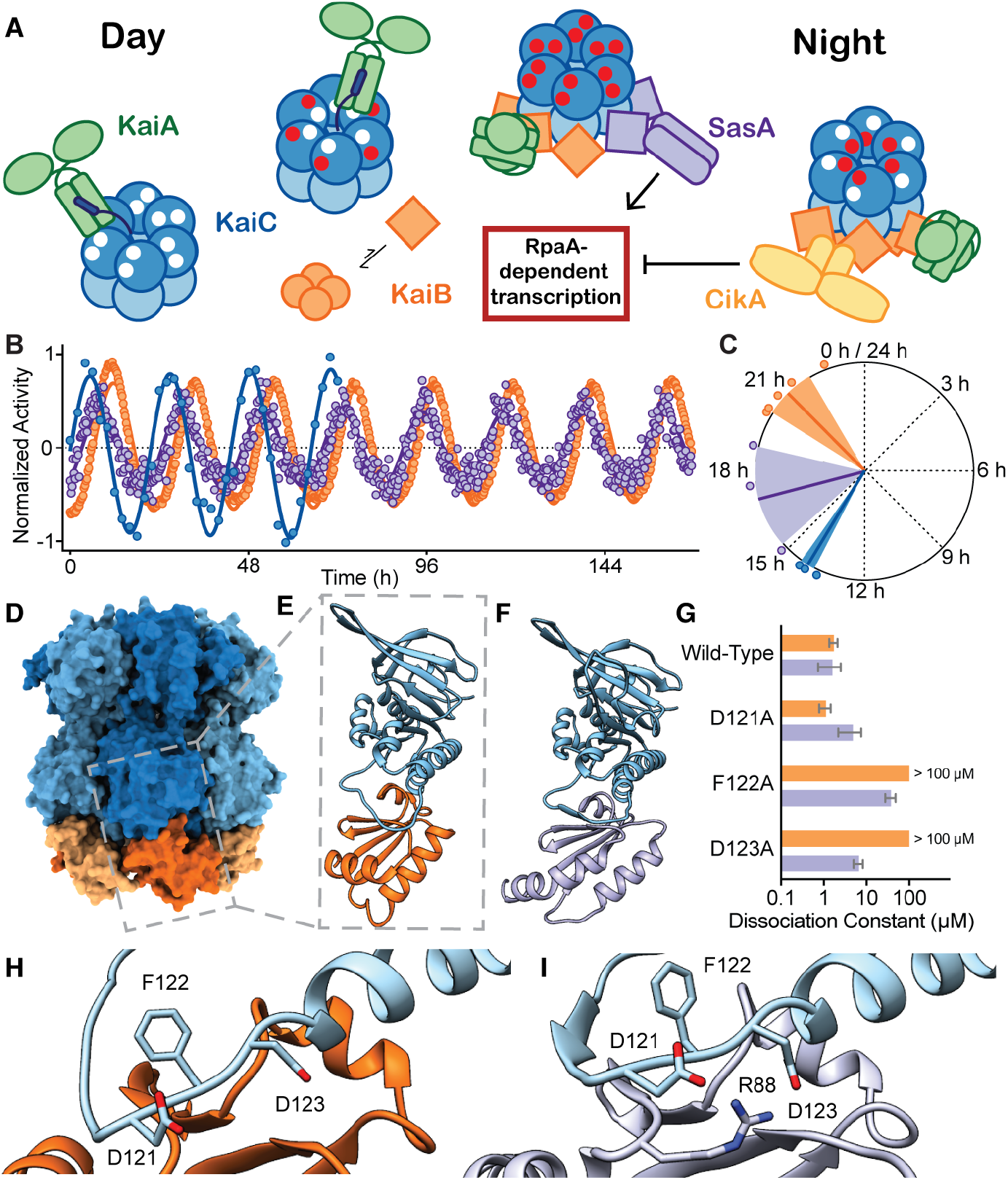
SasA and KaiB bind with sequential but overlapping phases to the same site on KaiC. **A**) Cartoon schematic of the cyanobacterial oscillator and associated output pathways. The KaiC phosphorylation rhythm (*18*) is depicted with white (unphosphorylated) or red (phosphorylated) circles. Ground-state KaiB tetramer (circles) interchanges with the fold-switched thioredoxin-like fold (squares). **B**) Normalized rhythms of KaiC phosphorylation (blue, measured by SDS-PAGE for 72 h) with KaiB or SasA_trx_ association with KaiC (orange or purple, respectively). Data points from individual traces (n=3) are overlaid in circles, and the mean (solid line) fitted to a sine function. See **Fig. S1** for raw data. **C**) Rayleigh phase diagram of KaiC phosphorylation and KaiB or SasA_trx_ binding for *in vitro* oscillations. Dark line, mean phase of KaiC binding or phosphorylation (circles represent n ≥ 3 assays; SD, width of the light wedge). Calculated phases are KaiB: 20.9 ± 0.9 h, SasA: 17.1 ± 1.7 h, and KaiC-P: 14.0 ± 0.4 h. **D**) The KaiB-KaiC hexamer (PDB:5JWQ). Subunits of KaiC are depicted in alternating light/dark blue, and fsKaiB monomers alternate yellow/orange. Gray box, position of the KaiC-CI domain bound to KaiB. **E**) The KaiC-CI domain-fsKaiB subcomplex (PDB:5JWO), with KaiC in blue, KaiB in orange. **F**) The KaiC-CI domain-SasA_trx_ subcomplex (PDB:6X61), with KaiC in blue, KaiB in orange. **G**) Equilibrium dissociation constants (K_D_) of KaiC-EE and mutants for KaiB or SasA_trx_ (mean ± SEM, n = 3). Where indicated, binding was too weak for curve-fitting and K_D_ is reported at > 100 μM. **H-I**) The interface of KaiC-CI with fsKaiB (**H**) or SasA_trx_ (**I**) with key residues highlighted.

To explore how the intrinsic structural similarity between fold-switched KaiB and the N-terminal thioredoxin-like domain of SasA might influence the cyanobacterial PTO, we took advantage of a new high-throughput fluorescence polarization-based post-translational oscillator (FP-PTO) assay that reports on complex formation with KaiC in real time (*17*). This allowed us to capture circadian rhythms of quaternary complex formation with KaiC under standard assay conditions for the PTO (*4*) in the presence of 50 nM fluorescently-labeled probes for KaiB or the thioredoxin-like domain of SasA, SasA_trx_, benchmarking the phase of these rhythms to the KaiC phosphorylation cycle (*18*) (**Fig. 1B** and **S1**). The phase of SasA_trx_ binding preceded that of KaiB by a few hours and was closer to the peak of KaiC phosphorylation (**Fig. 1C**), similar to that of full-length SasA (see accompanying paper by Chavan, A. et al.). Using KaiC phosphomimetics KaiC-EE and KaiC-EA, which are widely used to approximate the dusk-like pS,pT and nighttime-like pS,T states, we found that KaiB has similar affinity for both (**Fig. S2**). By contrast, full-length SasA has a higher preference for the earlier-occurring pS,pT state (*13*, *19*). Although the N-terminal domain of SasA is necessary and sufficient for binding to KaiC (*20*), avidity effects in the full-length dimer enhanced affinity for KaiC-EE by at least two orders of magnitude compared to the isolated, monomeric SasA_trx_ domain (*21*) (**Fig. S2**). One other contribution to the delayed phase of KaiB-KaiC binding could be its population shift from a highly stable tetrameric KaiB ground state to an unstable monomeric fold (*11*).

Six monomers of the active, thioredoxin-like fold of KaiB assemble onto the KaiC hexamer of *S. elongatus* (*8*) and the related thermophilic species, *T. elongatus* (*9*) to nucleate formation of the nighttime repressive state (**Fig. 1D**). A high-resolution crystal structure of the sub-complex comprising a single KaiC-CI domain and a fold switch-locked mutant of KaiB I88A (I87A in *S. elongatus*, referred to herein as “fsKaiB” for both) illustrates how KaiB docks onto the exposed B-loop of the CI domain (**Fig. 1E**). We solved a crystal structure of the SasA_trx_ domain bound to the KaiC-CI domain from *T. elongatus*, revealing that SasA binds the B-loop in a similar orientation to KaiB (**Fig. 1F** and **Table S3**). To interrogate the importance of this interface, we probed several KaiC residues in the B-loop by mutagenesis and found that substitutions at sites conserved between *S. elongatus* and *T. elongatus* (F122 and D123) decrease affinity for both KaiB and SasA (**Fig. 1G-I** and **S3**). We then examined how differences in the structures of the thioredoxin-like domains of KaiB and SasA, or their orientations on the B-loop might influence binding to adjacent subunits of the KaiC hexamer (**Fig. S3**). Modest changes in the length and orientation of the C-terminal helix between the SasA and KaiB thioredoxin-like folds could lead to steric clashes of SasA with a neighboring subunit. Consistent with this idea, saturation binding experiments showed that SasA_trx_ domains cannot fully occupy all six binding sites on the KaiC hexamer (**Fig. S3**). Moreover, unlike the highly cooperative binding observed for KaiB, SasA does not associate cooperatively with KaiC (*21*, *22*).

Prior studies of KaiB cooperativity demonstrated either one or six monomers of KaiB bound to the KaiC hexamer by native mass spectrometry (*21*, *22*). Given the similarity between the SasA-KaiC and KaiB-KaiC interactions, we wondered whether SasA could influence KaiB-KaiC interactions through heterotropic cooperativity. To test this idea, we performed equilibrium binding titrations of KaiC-EE with 50 nM fluorescently-labeled KaiB probe in the absence or presence of unlabeled SasA or the fsKaiB as a secondary titrant (referred to herein as an ‘additive’ to the KaiC-EE titration, **Fig. 2A, B**). While we could not easily assess the degree of homotropic cooperativity for KaiC-EE with the KaiB probe alone under these assay conditions, low concentrations (*e.g*., 50-100 nM) of SasA or fsKaiB significantly increased binding of the probe, demonstrating more efficient recruitment of KaiB to the KaiC hexamer. Higher concentrations of additives competed with the probe to delay its binding to the KaiC hexamer until the equivalence point where the concentration of KaiC-EE matches the additive. Interestingly, we observed a significant ‘overshoot’ of the typical KaiB probe anisotropy values at this point, indicative of positive heterotropic cooperativity (**Fig. 2A, B**). Binding of KaiB to the KaiC-CI monomer was not influenced by the addition of SasA or fsKaiB (**Fig. S4**), nor did SasA interact directly with KaiB in the absence of KaiC (**Fig. S4**), suggesting that these thioredoxin-like folds enhance the cooperative enhancement of KaiB only on the KaiC hexamer. While cooperativity operates at the KaiC hexamer level, a simplified two-site thermodynamic model was sufficient to account for the data using least-squares fitting. (**Fig. 2C** and **S4**). We defined the heterotropic ‘cooperativity index’ as the fold-increase in KaiB affinity given by the ratio of equilibrium constants K_1_/K_3_ (=K_2_/K_4_). Comparison of two-dimensional (2D) titration assays with SasA or fsKaiB to simulated data representing heterotropic or homotropic cooperative binding, respectively (**Fig. 2D**), to a model with no cooperativity (**Fig. 2E**), suggests that SasA and fsKaiB similarly influence cooperative recruitment of the KaiB probe on KaiC.

**Figure 2:**
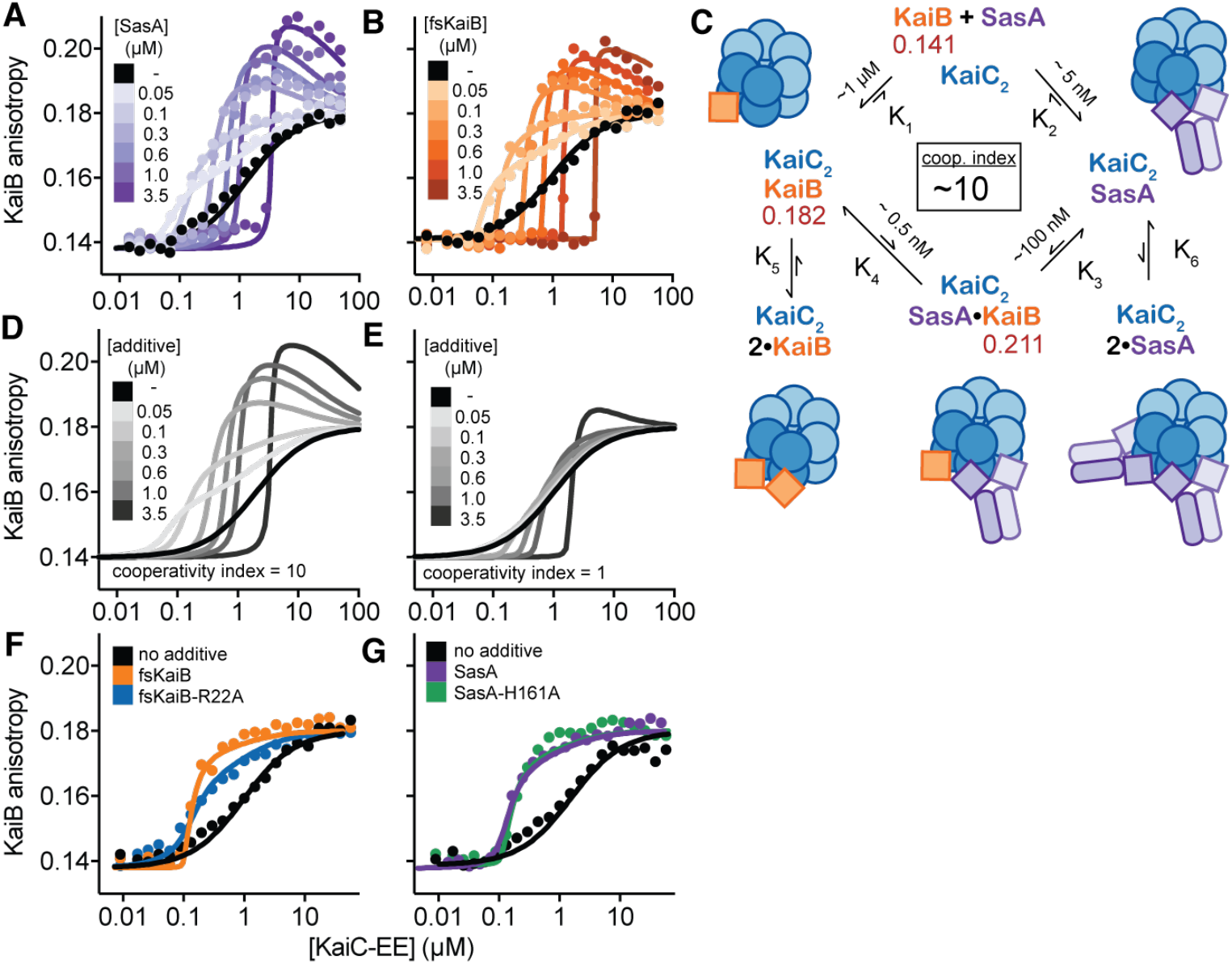
The thioredoxin-like fold of SasA and fsKaiB cooperatively recruits KaiB to the KaiC hexamer. **A-B**) Titrations of fluorescently-labeled KaiB with KaiC-EE in the absence (black) or presence of unlabeled full-length SasA (**A**, light to dark purple) or fsKaiB (**B**, light to dark orange). 2D titrations are shown from representative assays with global fitting to the thermodynamic model (see Fig. **S4**). **C**) Summary of two-site thermodynamic model for SasA heterocooperativity (darker shades) with the relevant higher-order species depicted in lighter colors. Mean equilibrium constants derived from least squares fitting of SasA 2D titration assays (n = 3) (**Data S3**); red, mean peak anisotropy values observed from global fitting of SasA 2D titration datasets. The cooperativity index is defined as the ratio K_1_/K_3_ = K_2_/K_4_. **D-E**) Simulated 2D titration data from thermodynamic model with a cooperativity index = 10 (**D**) or 1 (**E**). **F**) Titrations of fluorescently-labeled KaiB with KaiC-EE in the presence of 100 nM fsKaiB (orange) or fsKaiB-R22A (blue, full titration dataset in **Fig. S4). G**) Titrations of fluorescently-labeled KaiB with KaiC-EE in the presence of 100 nM SasA (purple) or SasA-H161A (green, full titration dataset in **Fig. S4**).

To explore the role of intersubunit interactions in KaiB cooperativity, we first turned to the KaiB-R22A mutant originally identified in *Anabaena* (*23*), which reduces the apparent affinity of KaiB for KaiC. This residue has been implicated in KaiB cooperativity (*21*) and is situated at the KaiB-KaiB interface in the KaiBC hexamer structure (*9*). When this mutant was tested in our cooperativity assay, we observed a decrease in the cooperative recruitment of the KaiB probe relative to fsKaiB (**Fig. 2G** and **S4**), demonstrating that inter-subunit KaiB-KaiB interactions play a role in this cooperative recruitment process. Moreover, although SasA kinase activity is required for its role in regulating circadian rhythms of transcriptional output *in vivo* (*20*), we found that the kinase-dead SasA mutant H161A is as effective as wild-type SasA in stimulating cooperative recruitment of KaiB to KaiC *in vitro* (**Fig. 2G**).

To see whether SasA’s role as a positive regulator of KaiBC association is important for its function *in vivo*, we set out to identify point mutations at potential cooperativity interfaces. We used crystal structures of SasA_trx_-CI and fsKaiB-C1 to model a SasA_trx_-KaiC-KaiB complex to examine potential SasA_trx_-KaiB interactions along the adjacent clockwise (CW) and counterclockwise (CCW) interfaces of the hexamer (**Fig. 3A**). Within this framework, we investigated a number of SasA mutations near the CW and CCW interface to see how they would influence cooperative recruitment in KaiB binding assays (**Fig. S5**). Using analysis of 2D titration data with the mutants, we found a decrease in the cooperativity indices for the individual mutants H28A (CCW interface) or Q94A (CW interface) that was further decreased in the H28A/Q94A double mutant (**Fig. 3B**). The H28A/Q94A displayed a striking lack of heterocooperativity in the 2D titration assay (**Fig. 3C** compared to simulated data in **2E**), demonstrating that SasA-KaiB interactions at both interfaces are important for cooperative recruitment.

**Figure 3:**
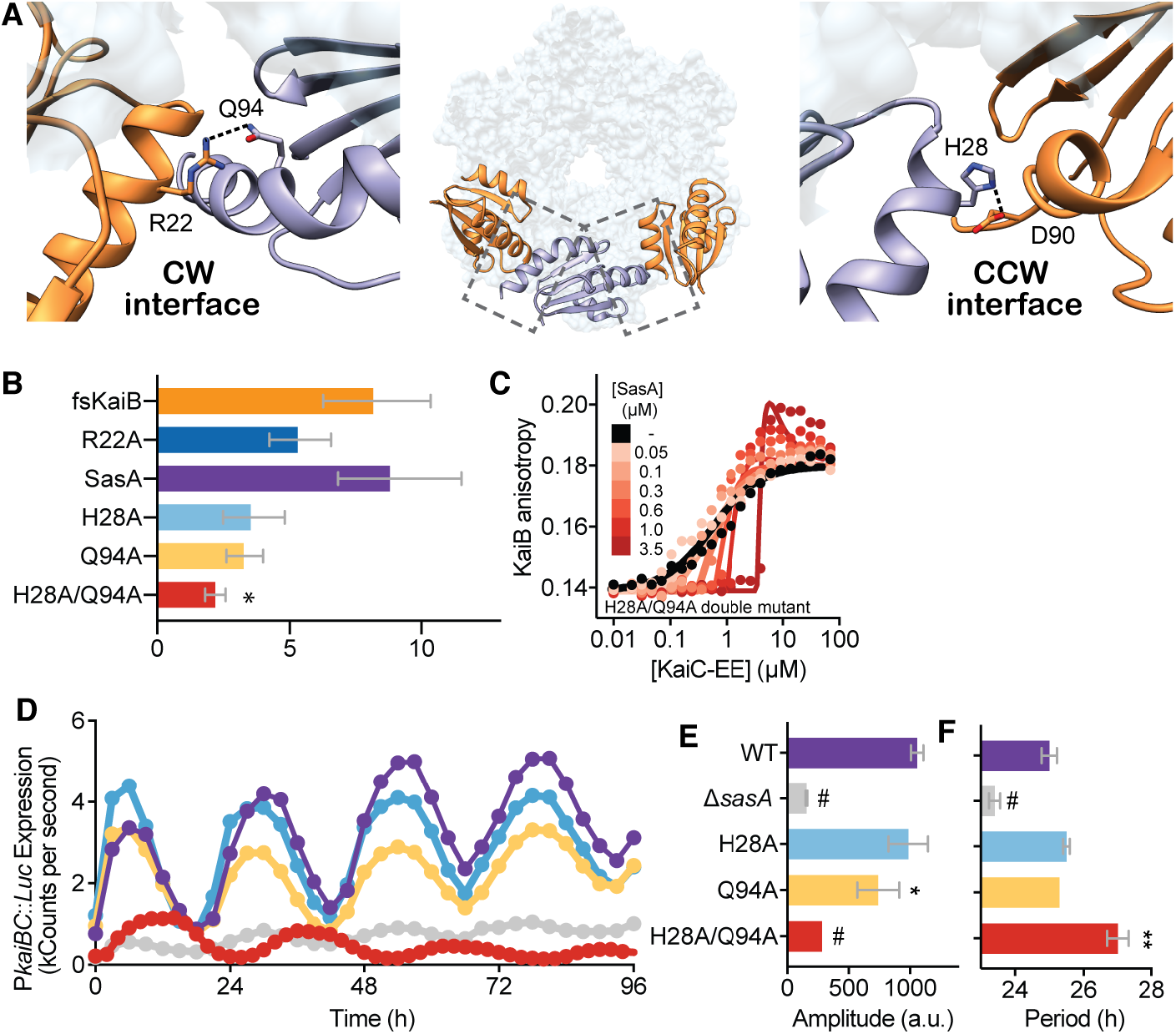
SasA-KaiB interactions mediate cooperative recruitment of KaiB *in vitro* and sustain robust circadian rhythms *in vivo*. **A**) Structural model of KaiB:SasA interactions on KaiC from KaiC-CI subcomplexes with fsKaiB (orange, PDB:5JWO) or SasA_trx_ (purple, PDB:6X61) modeled onto adjacent subunits of KaiC (light blue) of the KaiBC hexamer (PDB:5JWQ). Key residues at clockwise (CW) and counterclockwise (CCW) interfaces are modeled as *S. elongatus* variants based on the alignment in **Fig. S5**. Dotted lines, polar contacts predicted by hybrid structural model. **B**) Cooperativity indices for KaiB and SasA mutants from thermodynamic model. Bars represent median and 95% confidence intervals from Monte Carlo analysis from n = 2-3 full 2D datasets. See Supplemental Methods for more details. Statistical analysis was performed on variance between median Monte Carlo values between replicates (*, *P* < 0.05) **C**) Representative 2D titration of fluorescently-labeled KaiB with KaiC-EE in the absence (black) or presence of SasA-H28A/Q94A (light to dark red). **D**) Representative timecourse of bioluminescence driven by P*kaiBC* from *S. elongatus* cultures entrained under 12-h LD cycles for 48-hr and subsequently allowed to free run in LL. Raw luminescence curves were fit to a sine function to extract amplitude (**E**) and free running circadian period (**F**) for each strain. Error bars depict standard deviation among replicate cultures (n = 6-12); when error bars are not visible, they were smaller than could be depicted. Symbols indicate significance (**, *P* < 0.01; #, *P* < 0.0001) from one-way ANOVA of mutants relative to wild-type *S. elongatus*.

To see if the SasA-induced heterocooperativity we observed *in vitro* influences circadian rhythms *in vivo*, we introduced these substitutions into SasA using CRISPR/Cas12a and monitored bioluminescence from a P*kaiBC* luciferase reporter in constant light (LL) after synchronization in 12-h light:12-h dark cycles (**Fig. 3D** and **S6**). Modest effects on amplitude were seen with the individual H28 or Q94 alanine substitutions on their own, but when these were combined in the H28A/Q94A mutant, the amplitude of circadian rhythms was decreased to an extent that was similar to the SasA knockout strain (*ΔsasA*, **Fig. 3E**). Furthermore, we observed a 2-h increase in period in the double-mutant strain (**Fig. 3F**), suggesting that heterocooperative interactions at the CW and CCW interfaces are redundant, but crucial for circadian timekeeping *in vivo*.

SasA was originally described as an amplifier of circadian rhythms, critical for maintaining robust rhythmicity by controlling rhythmic transcription of the *kaiBC* cluster (*20*). To test whether this amplitude-enhancing effect might be due in part to SasA regulation of KaiB binding to KaiC, we turned to the FP-PTO assay to explore the effects of SasA on robustness of the core circadian oscillator in the absence of transcription-translation feedback that occurs *in vivo* (*1*). First, we measured the dependency of this oscillator assay on the concentrations of KaiA and KaiB, comparing data from the FP-PTO to a prior study using rhythms of KaiC phosphorylation to find a striking convergence in our results (**Fig. 4A, B**) (*24*). Because the FP-PTO is monitored non-invasively in 384-well format, we could follow the oscillator under different conditions for many days longer than is practical for the KaiC phosphorylation assay that is analyzed by SDS-PAGE. Generally, oscillator amplitude and period were highly sensitive to KaiA concentrations, and while there was a clear requirement for a minimal concentration of KaiB, the period was quite robust to excess KaiB relative to KaiC (**Fig. 4C-E**). Although the oscillator functioned at low amplitude for several days with a normal circadian period with one-half stoichiometry of KaiB relative to KaiC (i.e., 1.75 μM KaiB for 3.5 μM KaiC) (*24*), these rhythms rapidly damped (**Fig. 4B**). KaiB is present in only modest excess over KaiC *in vivo* (*25*, *26*), which suggests that the ability of KaiC to cooperatively recruit and bind six KaiB monomers is critical to sequester KaiA stably during negative feedback in the PTO and is thus a determinant of clock robustness. Consistent with this idea, the amplitude of the PTO decreased as the concentration of KaiA approached or exceeded that of KaiC (**Fig. 4A, C** and (*13*, *24*)).

**Figure 4:**
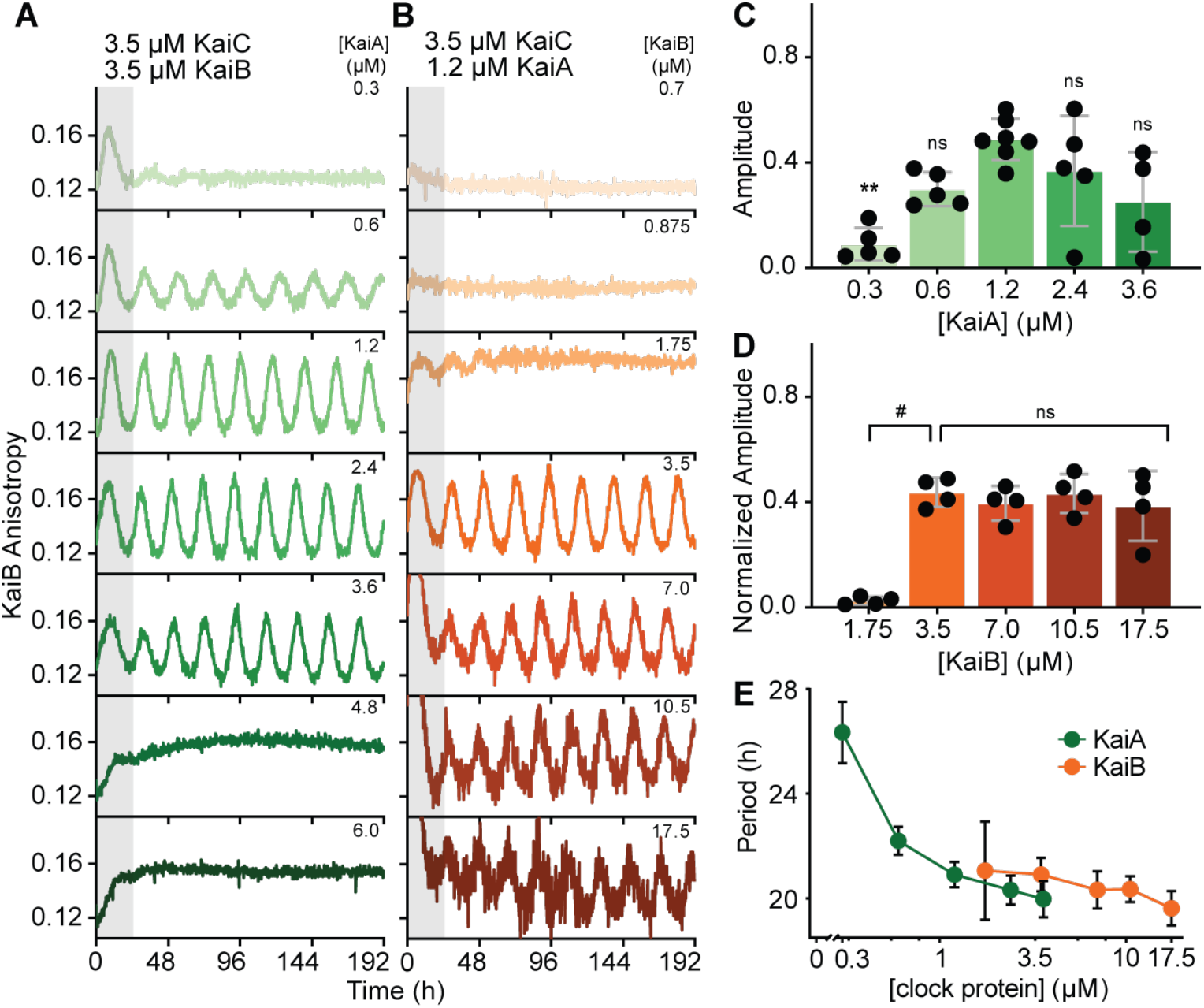
Amplitude and period of the oscillator depend in different ways on the concentrations of KaiA and KaiB. **A)** The FP-PTO *in vitro* with standard conditions of 3.5 μM KaiC, 3.5 μM KaiB (including 50 nM fluorescently-labeled KaiB as a probe) and titrations of KaiA from 0.3-6.0 μM. Representative assay from n =3 shown; the first 24-hr period after release into constant conditions is marked in gray. **B**) The FP-PTO with standard conditions of 3.5 μM KaiC, 1.2 μM KaiA and titrations of KaiB from 0.7-17.5 μM (including 50 nM fluorescently-labeled KaiB as a probe). **C-D**) Amplitude analysis of KaiA titrations (**C**, data from panel **A**) or KaiB titrations (**D**, data from panel **B**). Amplitudes for KaiB titrations were normalized to account for increasing ratio of unlabeled KaiB to fluorescently-labeled KaiB. Reaction conditions that could not be fit by FFT-NLLS analysis, and therefore did not oscillate, were not included. Data are shown as mean ± SD (n ≥ 3). One-way ANOVA with Dunnett’s multiple comparisons test was used for comparison between groups: ns, non-significant; *, *P* < 0.05; **, *P* < 0.01; ***, *P* < 0.001; #, *P* < 0.0001. **E**) Period of the FP-PTO plotted against KaiA (green) or KaiB (orange) concentrations. Data are shown as mean ± SD (n ≥ 3).

Because of its ability to enhance formation of the KaiBC complex, SasA has the potential to improve robustness of the PTO under limiting concentrations of KaiB by recruiting a stable, stoichiometric complement of KaiB to KaiC hexamers to sequester KaiA. To explore how SasA influences KaiB binding in the context of an oscillator, we set up FP-PTO assays under typical *in vitro* oscillator conditions (*4*) as well as under limiting concentrations of KaiB relative to KaiC. Low anisotropy levels for labeled KaiB under the limiting condition of only 0.875 μM KaiB (**Fig. 5A**, left column) demonstrated incomplete KaiB binding to the KaiC hexamer. The addition of full-length SasA up to 1 μM increased KaiB anisotropy levels, demonstrating that SasA enhanced KaiB binding to KaiC in the presence of KaiA, even without a functional oscillator. However, as the concentration of SasA approached or exceeded that of KaiC, anisotropy values for KaiB dropped, indicating that SasA outcompeted KaiB for binding to the KaiC hexamer as observed before (*12*, *27*, *28*). When KaiB was present at one-half stoichiometry with respect to KaiC (1.75 μM KaiB for 3.5 μM KaiC), the addition of SasA restored robust oscillations to the FP-PTO (**Fig. 5A**, middle column). This enhancement appears to be almost catalytic, as only 100 nM SasA was sufficient to rescue damping (**Fig. 5B**) and restore amplitude (**Fig. 5C**) to wildtype levels. Increasing SasA concentrations to 1 μM under these conditions did not influence amplitude any further, while higher concentrations of SasA outcompeted KaiB binding to attenuate oscillations in the FP-PTO (**Fig. 5C**).

**Figure 5:**
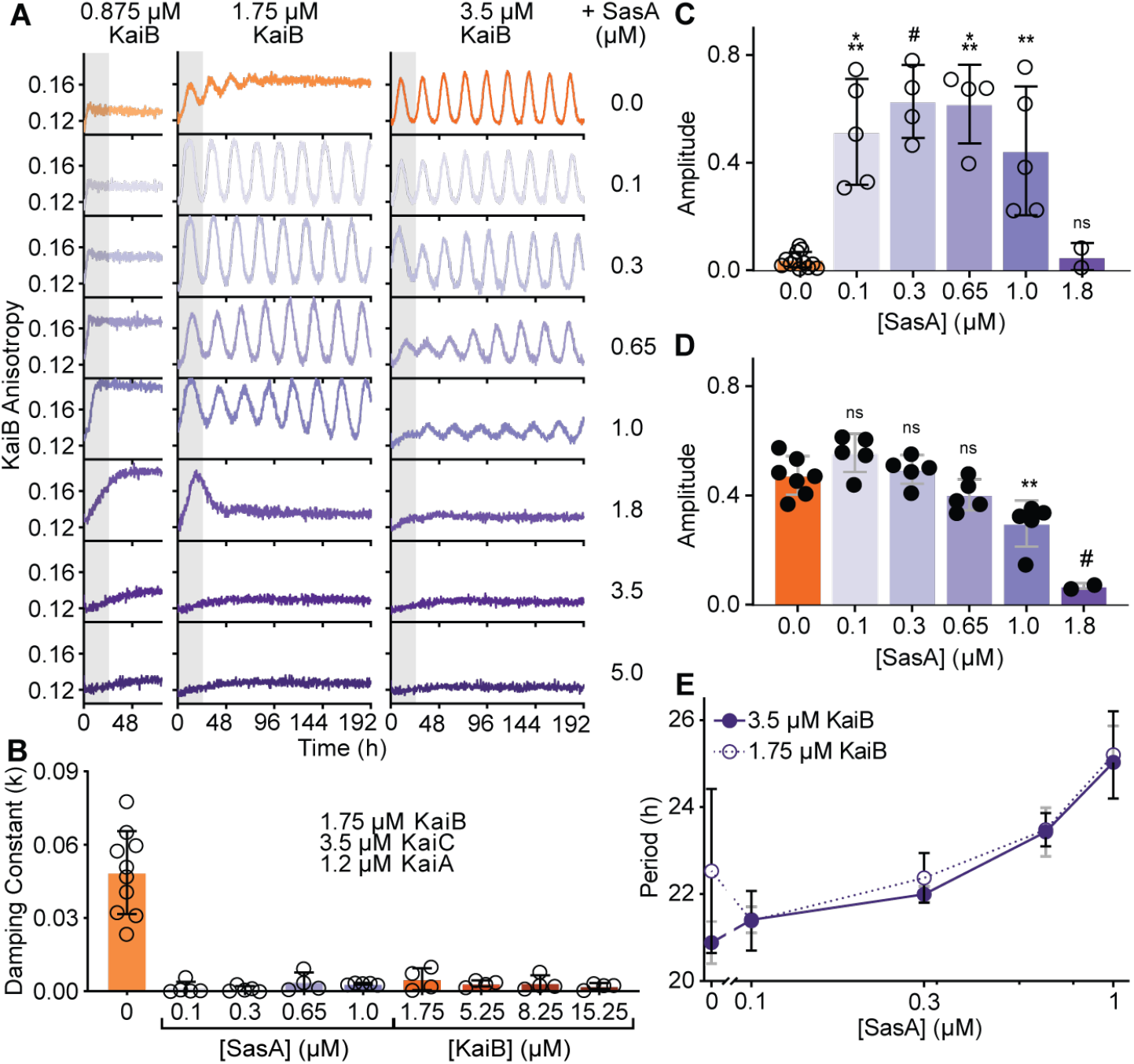
SasA dynamically influences robustness of the post-translational oscillator *in vitro*. **A**) The FP-PTO with 3.5 μM KaiC, 1.2 μM KaiA, and the indicated concentrations of KaiB (including 50 nM fluorescently-labeled KaiB as a probe) with titrations of full-length SasA 0.1-5.0 μM. Representative assay from n ≥ 3 shown; the first 24-hr period after release into constant conditions is marked in gray. **B**) Damping constants of the FP-PTO with 1.75 μM KaiB (including 50 nM fluorescently-labeled KaiB as a probe) in the presence of added SasA (purple) or KaiB (orange). **C-D**) Amplitude of the FP-PTO prepared with 1.75 μM KaiB (**C**, open circles) or 3.5 μM KaiB (**D**, closed circles) in the presence of SasA (light to dark purple). **E**) Period of the FP-PTO prepared with 1.75 μM KaiB (open circles) or 3.5 μM KaiB (closed circles) with SasA (light to dark purple). Reaction conditions that could not be fit by FFT-NLLS analysis, and therefore did not oscillate, were not included. Damping constant, period, and amplitude data are representative of four or more independent experiments, are presented individually and as the mean ± SD. One-way ANOVA with Dunnett’s multiple comparison test was used for comparison between groups: ns, non-significant; *, *P* < 0.05; **, *P* < 0.01; ***, *P* < 0.001; #, *P* < 0.0001.

Under standard oscillator conditions with stoichiometric KaiB and KaiC, the presence of SasA up to 0.65 μM did not markedly influence the oscillator (**Fig. 5A**, right column), whereas at increased concentrations of SasA, oscillator amplitude diminished until KaiB was completely outcompeted for KaiC binding (**Fig. 5B, D**). This suggests that the competitive effects of SasA may be enhanced as the concentration of KaiB equals KaiC. Moreover, the sharp drop in amplitude of the FP-PTO suggests that the functional switch from cooperativity to competition occurs around ~1 μM SasA depending on the exact oscillator conditions; interestingly, this ultrasensitive setpoint is close to the concentration of SasA *in vivo* estimated by quantitative western blotting (0.58 ± 0.07 μM from (*29*)). In all cases where conditions were sufficient to generate a stable PTO, the period was lengthened similarly by addition of SasA (**Fig. 5E**). Therefore, in addition to regulating rhythms of transcriptional output from the Kai-based PTO (*20*), SasA also works to directly modulate KaiB association with KaiC to control formation of the nighttime repressive state and robustness of the PTO itself.

The other circadian output kinase, CikA, also contributes directly to robustness of the PTO by enhancing rhythms under limiting concentrations of KaiA (*16*). Using the FP-PTO assay, we observed that addition of CikA moderately enhanced low amplitude rhythms under limiting concentrations of KaiA while shortening the period (**Fig. S7**). The pseudo-receiver (PsR) domain of CikA binds to the same site on KaiB in the nighttime complex as KaiA does (*9*), and their competition *in vivo* is made evident by the phenotypes of a *kaiA* deletion mutant (*30*). By competing KaiA out of the repressive complex, CikA promotes the activating potential of KaiA to stimulate KaiC phosphorylation (*6*, *16*). Either the isolated PsR domain or full-length CikA caused similar concentration-dependent decreases in period length in the FP-PTO assay, while the isolated thioredoxin-like domain of SasA was much less effective at lengthening the period compared to full-length SasA, likely due to avidity effects (**Fig. S2** and **S8**).

The FP-PTO assay implemented here has uncovered roles for the output kinases as accessory components of the oscillator, expanding its functionality beyond the narrow concentrations and ratios that are tolerated in the traditional *in vitro* oscillator (*24*). This extended oscillator reveals how the clock maintains consistency *in vivo* throughout rhythmic changes in oscillator components that occur as part of transcription-translation feedback (*26*, *31*) and protein turnover (*32*, *33*), as well as providing an experimental platform for integrating the oscillator with the upstream and downstream components with which it interacts (see manuscript by Chavan, A. et al.).

## Acknowledgments

We thank staff at the 23-ID-D beamline of the Advanced Photon Source, Argonne National Laboratory for their help with data collection. The Advanced Photon Source (contract DE-AC02-06CH11357) is supported by the U.S. Department of Energy. Molecular graphics and analyses performed with UCSF Chimera and ChimeraX, developed by the Resource for Biocomputing, Visualization, and Informatics at the University of California, San Francisco, with support from National Institutes of Health P41-GM103311(Chimera) and R01-GM129325 and the Office of Cyber Infrastructure and Computational Biology, National Institute of Allergy and Infectious Diseases (ChimeraX). We thank Archana G. Chavan, Michael Rust, Lu Hong, Shahar Sukenik, Maria Zoghbi for useful discussions.

## Author contributions

Conceptualization, J.H., J.A.S., C.L.P., and A.L.; Methodology, J.H., J.A.S., and C.R.B.; Investigation, J.H., J.A.S., J.G.P., C.S., D.C.E., R.K.S., and S.T.; Validation, J.H., J.A.S., and S.T.; Formal Analysis, J.H., J.A.S., C.R.B., and P.C.; Resources, S.S.G; Data Curation, J.H. and J.A.S.; Writing – Original Draft, J.H., J.A.S., and C.L.P.; Writing – Reviewing and Editing, J.H., J.A.S., D.C.E., C.R.B., S.S.G., C.L.P., and A.L.; Funding Acquisition, J.H., J.P., P.C., S.S.G., C.L.P., and A.L.; Supervision, S.S.G., C.L.P., and A.L.

## Data and materials availability

All data is available in the main text or Supplementary Materials.

## Supplementary Materials

Materials and Methods

Figures S1-S8

Tables S1-S10

External Data S1-S6

References (1-27)

