## Supplementary Material for "Structural mimicry confers robustness in the cyanobacterial circadian clock"

**This PDF file includes:**

Materials and Methods
Figs. S1 to S8
Tables S1 to S9
Captions for Data S1 to S6
Supplemental References 1-27

**Other Supplementary Materials for this manuscript include the following:**

Data S1 to S6

### Materials and Methods

Cloning of constructs

PCR-mediated mutagenesis was performed on the pET-28b vector utilizing Nde I/Hind III cut sites as described previously (1). Full-length *sasA* was cloned into the pHis-Gβ1 parallel expression vector using EcoRI/NotI cut sites (2,3). All constructs are listed in **Table S1**. Point mutations were introduced using long-range PCR with overlapping primers (4).

Protein expression and purification

Protein expression for most constructs was carried out in BL21(DE3) *E. coli* (Novagen) and purified by Ni-NTA affinity chromatography and size-exclusion chromatography as described previously (1). All buffers for KaiC purification included 1 mM ATP and 5 mM $MgCl_2$ added as described Chang et al. (1). Fluorescence labeling of KaiB, $SasA_{trx}$, and $CikA_{psr}$ constructs was done with 6-iodoacetamidofluorescein (6-IAF, Invitrogen) as described previously (5).

After purification of His-Gβ1-tagged full-length SasA constructs over Ni-NTA resin using the standard protocol described in Chang et al. (1), the protein was cleaved overnight at 4°C with TEV protease a final concentration of 0.1 mg/mL. SasA was subsequently resolved from the His-Gβ1 tag on a Sephadex 200 size-exclusion column (GE Healthcare) equilibrated with 20 mM Tris pH 7.4, 150 mM NaCl.

Fluorescence Polarization - Post Translational Oscillator (FP-PTO) assays

Fluorescence anisotropy oscillator data were collected *in vitro* on either a CLARIOstar Plus (BMG) or Spark 10M (TECAN) microplate reader. The fluorescein channel was used for all data collection ($\lambda_{excitation}$, 490 ± 5 nm; $\lambda_{emission}$, 520 ± 5 nm). Fluorescence anisotropy was monitored from each well in the 384-well plate and recorded every 15 minutes, with time zero representing 3-5 minutes following the addition of KaiC to oscillation reactions. See **Table S2** for specific experimental conditions for each run.

FP-PTO data quantification and statistical analyses

Fluorescence anisotropy readings from FP-PTO assays were collected in MARS Data Analysis Software or SparkControl Software for experiments run on the BMG CLARIOstar Plus or TECAN Spark 10M, respectively. All data were analyzed in the online BioDare suite by FFT-NLLS (6,7) (https://biodare2.ed.ac.uk/welcome). Prior to analysis, fluorescence anisotropy rhythms were baseline detrended and normalized to [-1, 1] with mean of zero. The first 12-h of data were disregarded for quantification of period, amplitude, and phase. Period (**Table S4**) and amplitude (**Table S5**) analysis in **Fig. 4, 5, S7,** and **S8** were assessed by ordinary one-way ANOVA with Dunnett's multiple comparison tests in Prism 8 (Graph Pad). The significance values and the number of independent experiments for each experimental group are reported in the corresponding figure legends.

Comparisons of the effects of added SasA or CikA on period (**Table S6**) and amplitude (**Table S7**) under different concentrations of the core clock proteins KaiA and KaiB presented in **Table S2** were determined by ordinary two-way ANOVA with Dunnett's multiple comparison tests in Prism 8 (GraphPad). Normalized fluorescence anisotropy rhythms were plotted in conjunction with nonlinear regression least squares cosinor fit in Prism 8 in **Fig. 1**, while raw fluorescence anisotropy data were plotted with Origin Student 2019b (Origin Lab) in **Fig. 4, 5, S7,** and **S8**.

Raw anisotropy data were baseline corrected using an unbound labeled-KaiB reaction well as a reference in each independent experiment. The phase diagram in **Fig. 1C** was prepared using phase values resulting from analysis by BioDare2 (6,7) as described above.

The damping constant, k, was determined in Prism 8 (GraphPad) using a nonlinear least squares regression cosinor fit:

$$y = m * x + amplitude * e^{-kx} * \cos\left[\left(2\pi * \frac{x}{period}\right) + phase\right]$$

where y is the signal, x the corresponding time, amplitude is the height of the peak of the waveform above the trend line, k is the decay constant (such that 1/k is the half-life), period is the time taken for a complete cycle to occur and phase is the shift in x relative to a cosinor wave.

KaiC phosphorylation assay

KaiC phosphorylation assays were performed *in vitro* as previously described (8). Densitometry was used to quantify KaiC phosphorylation using ImageJ FIJI (NIH). Quantification of phases for phosphorylation densitometry rhythms was assessed in the online BioDare2 suite (6,7) as described above.

Crystallization of monomeric *T. elongatus* C1 domain in complex with *T. elongatus* $SasA_{trx}$

A monomeric mutant of *T. elongatus* KaiC-CI domain (see **Table S1** for details) was incubated at 250 µM with an excess of *T. elongatus* $SasA_{trx}$ (460 µM) overnight in 20 mM Tris pH 7.0, 150 mM NaCl, 5 mM DTT, 1 mM $MgCl_2$ and 1 mM ATP at room temperature. The complex was subsequently purified by size-exclusion chromatography on a Sephadex 70 column (GE Healthcare) equilibrated in the same buffer, but with $MgCl_2$ and ATP concentrations reduced to 0.5 mM. The complex was mixed in a 1:1 ratio to a final concentration of 10.8 mg/mL with the crystallization buffer containing 1.26 M $NaH_2PO4$, 0.54 M $K_2HPO_4$ (pH unadjusted, total $PO_4$ concentration 1.8 M), 0.1 M Glycine (added from a 1 M solution adjusted to pH 10.5) and 0.2 M $Li_2(SO_4)_2$. Crystals formed over 10 days at 22 °C using the hanging drop method. The flat, plate-like crystals were then frozen in liquid nitrogen after soaking in cryoprotectant composed of the crystallization buffer plus 20% (v/v) glycerol.

Structure determination and refinement

Single crystal diffraction data were collected with a wavelength of 1 Å on the 23-ID-D X-ray source at the Advanced Photon Source at the Argonne National Laboratory. Data were processed and scaled using MOSFLM (9) and Aimless (10). Phases were solved by molecular replacement with the structure of *T. elongatus* KaiC-CI monomer in complex with fsKaiB (PDB 5JWO) using Phaser (11). Refinement and model building were performed using Phenix (12) and Coot (13). See **Table S3** for crystal and refinement statistics. Structural figures were made using UCSF Chimera (14,15) and ChimeraX (16).

Equilibrium binding assays

Binding titrations were performed in 20 mM Tris pH 7.4, 150 mM NaCl, 1 mM ATP, 1 mM $MgCl_2$ and 0.1 % (v/v) Tween-20. Fluorescein-labeled KaiB or $SasA_{trx}$ probes were present at 50 nM, while the titrant was diluted serially in 1/3-fold increments. Serial dilutions were performed in a 384-well plate before sealing with tape and incubating at room temperature overnight (9-15 h). Fluorescence polarization anisotropy measurements were subsequently collected on a

Synergy2 plate reader (BioTek). Replicate measurements were collected and averaged for each well (20). For 2D titration assays looking at the effect of an additive on KaiB binding to KaiC, fsKaiB or SasA additives were included in both KaiC and diluent buffer to maintain a constant concentration. Diluent was added to the 384-well plate using a single channel pipettor, and additives were mixed into the KaiC stock last and diluted within 10 minutes. See thermodynamic modeling of binding equilibria below for more information.

Thermodynamic modeling of binding equilibria

The fluorescence anisotropy titrations outlined above involve cooperative and competitive reactions and span a wide range of concentrations, such that free ligand concentrations cannot be approximated by the total added concentration. Consequently, these data cannot be analyzed by fitting to standard analytical equations (17). Fitting the data to the profiles simulated by a model avoids this problem but introduces others in terms of the complexity of the model that is required for the fit. Initially, we attempted to fit to a general hexameric model for KaiC but found there were too many parameters to reach convergence. When simplified to a dimer model, the fits were reasonably robust and showed no systematic deviations. Nevertheless, such a simplified model required positive heterotropic or homotropic cooperativity between KaiB and additives such as SasA and fsKaiB, respectively, as well as competition between these additives. Thus, a dimer model captures the essence of the interaction, although how this relates in detail to cooperativity within the KaiC hexamer remains in question.

Least squares fitting analysis to models was performed using DynaFit (BioKin) (18). Scripts used for analysis are available in supplemental data (**Data 1** and **2**). Statistical analysis was performed using the Monte Carlo routine. Cooperativity indices (described by $K_1/K_3 = K_2/K_4$, see **Data S3**) were calculated for each simulation ($n = 1000$) and median and 95% confidence intervals taken as ranks 500, 25 and 975 (respectively) in the $n = 1000$ simulation. Where replicate measurements are reported, the values of median or 95% confidence boundaries were averaged amongst the replicates.

In order to reduce the number of parameters of the fit, the binding of KaiB alone was initially modeled without any homotropic cooperativity by assigning $K_5 = 4*K_1$, as is appropriate for the macroscopic equilibrium constants for two-site independent binding. When $K_5$ was floated, a slightly improved fit was obtained with a returned $K_5 < K_1$, indicative of homotropic cooperativity, but $K_5$ was not robustly defined. The value of the heterotropic cooperativity index in the presence of additives increased when $K_5$ was floated, however, we report fits where $K_5$ was defined as $4*K_1$ for simplicity, which gives a minimal estimate of the heterotropic cooperativity index. The input and output files from the least-squares fitting and Monte Carlo analysis are available as **Data S3**. Input KaiC dimer concentrations are given in Data S3, while KaiC concentrations given in terms of total monomer elsewhere.

Triplicate 2D titrations were collected with SasA to optimize the analysis, and showed some variability that was ameliorated by floating additive concentrations at the 3 highest additive concentrations. The averages from these fits were used for SasA and KaiB variant 2D titration datasets when analyzing 300 nM data, where additive concentrations were also allowed to float. Little variation was seen in the experimental anisotropy values determined for fluorescently-labeled KaiB alone or the final peak values for the KaiB-KaiC complex, though the average peak experimental anisotropy values of putative ternary complexes seeded by heterocooperativity differed modestly between the SasA and fsKaiB variants (KaiB peak anisotropy = 0.211 for SasA or 0.205 for fsKaiB).

Size-exclusion chromatography-multiangle light-scattering (SEC-MALS) assays
SEC-MALS assays were performed at room temperature using a silica-based size-exclusion column (particle size 5 μm, pore size 500 Angstrom, 4.6 mm ID, Cat. No. WTC-050N5, Wyatt Technologies) to resolve the oligomeric state of SasA. 20 μL injections of full-length SasA at 1.5 mg/mL were made using an Agilent G1311A quaternary pump and manual injector (Rheodyne), run over the silica-based column, and analyzed by a T-rEX refractometer and miniDAWN TREOS II static multiangle light scattering instrument (Wyatt Technologies) directly after the column. Analysis of absolute molecular weight was carried out using Astra 6.0 software (Wyatt Technologies).

Generation of SasA mutants in *S. elongatus*
Markerless point mutations were introduced in *sasA* of *Synechococcus elongatus* PCC 7942 by CRISPR/Cas12a engineering as previously described (19). Plasmids and the primers used in vector construction and sequence verification are listed in **Table S8**. Briefly, oligos with complementarity to the guide RNA (gRNA) recognition site were annealed and cloned into AarI-cut pSL2680 (Addgene Plasmid #85581). Clones of pSL2680 that carry the appropriate gRNA insert were isolated and plasmid sequences were verified. Upstream and downstream homologous repair templates that encode the point mutation(s) of interest were amplified by PCR and assembled (GeneArt Seamless Assembly, Thermo Fisher) into KpnI-cut constructs that contain the respective gRNAs. Recovered plasmids were checked for accuracy by Sanger sequencing prior to editing in *S. elongatus*.

The RSF1010-based editing constructs were electroporated into *E. coli* AM1359 that contain conjugal helper plasmids (pRL623 and pRL443) as previously described (20-24). The resulting *E. coli* strains were grown overnight in LB containing ampicillin (100 μg/ml), chloramphenicol (17 μg/ml) and kanamycin (50 μg/ml). Cells from a 1 ml aliquot were washed three times with fresh LB and resuspended in a final volume of 100 μl LB, then mixed with 100 μl of an *S. elongatus* clock-reporter strain (AMC541) concentrated down from 2 ml of a dense culture ($OD_{750}$ = ~0.6). The mixed culture was plated to solid BG-11 medium containing 5% LB (v/v) and incubated at 30˚C under 30 μmol photons $m^{-2}$ $s^{-1}$ (μE) illumination for 24 hours. Plates were then underlaid with kanamycin (5 μg/ml final concentration) to select for the editing plasmid. *S. elongatus* colonies that emerged after 8-10 days at 30˚C and 100 μE light were serially patched three times to BG-11 containing kanamycin to maintain the editing plasmid long enough to complete segregation of the mutant allele in all copies of the chromosome. After editing, *sasA* was amplified by colony PCR using primers that anneal outside of the homologous repair region and the resulting PCR product was submitted for Sanger sequencing to confirm segregation of the point mutation(s) of interest.

Bioluminescence monitoring of *S. elongatus* circadian rhythms
The impact of *sasA* mutations on output from the KaiABC oscillator *in vivo* was measured using a firefly luciferase reporter driven by the *kaiBC* promoter ($P_{kaiBC}$::*luc*) as previously described (25). Mutant *sasA* strains, along with positive and negative clock-output controls, were grown in

BG-11 medium, diluted to $OD_{750}$ = 0.2 and arrayed in 96-well plates containing solid BG-11 medium and 10 µl of 5 mM D-luciferin. Plates were covered by a gas permeable seal and incubated in a light-dark chamber at 30˚C for 48 hours, with 12 hour intervals of 120 µE light and darkness. Following release into constant light at the end of 48 hours, plates were transferred to a lighted stacker (40 µE light) attached to a Tecan Infinite M200 Pro and bioluminescence was monitored every 2-3 hours. The raw bioluminescence data were plotted as a function of time (GraphPad Prism 8) and processed using BioDare2 to determine period and amplitude for each set of replicates (7). All strains used in this study are listed in **Table S9**.

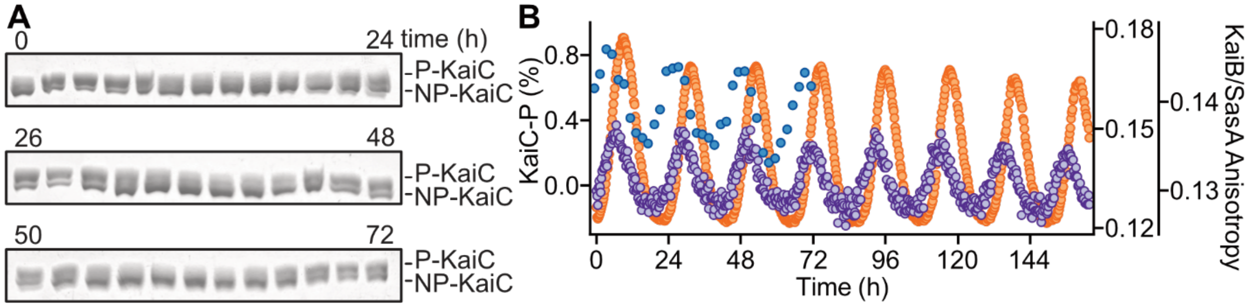

**Figure S1. KaiC phosphorylation and FP-PTO of fluorescently-labeled KaiB and SasA.**

**A**) KaiC phosphorylation rhythms under the standard oscillator concentrations of 3.5 μM KaiC, 3.5 μM KaiB, and 1.2 μM KaiA *in vitro*. P-KaiC, phosphorylated KaiC; NP-KaiC, non-phosphorylated KaiC. **B**) Overlay of KaiC phosphorylation (KaiC-P) rhythm (blue) with unnormalized (raw) FP values for fluorescently-labeled KaiB probe (orange, inner right y-axis) or fluorescently-labeled $SasA_{trx}$ probe (purple, outer right y-axis) from distinct FP-PTO assays set up under standard *in vitro* oscillator conditions and plotted against the start of incubation, with time zero representing the moment KaiC was added to the reactions(see **Fig. 1B** for normalized data). The smaller change in anisotropy exhibited by the $SasA_{trx}$ probe may reflect more transient association with KaiC.

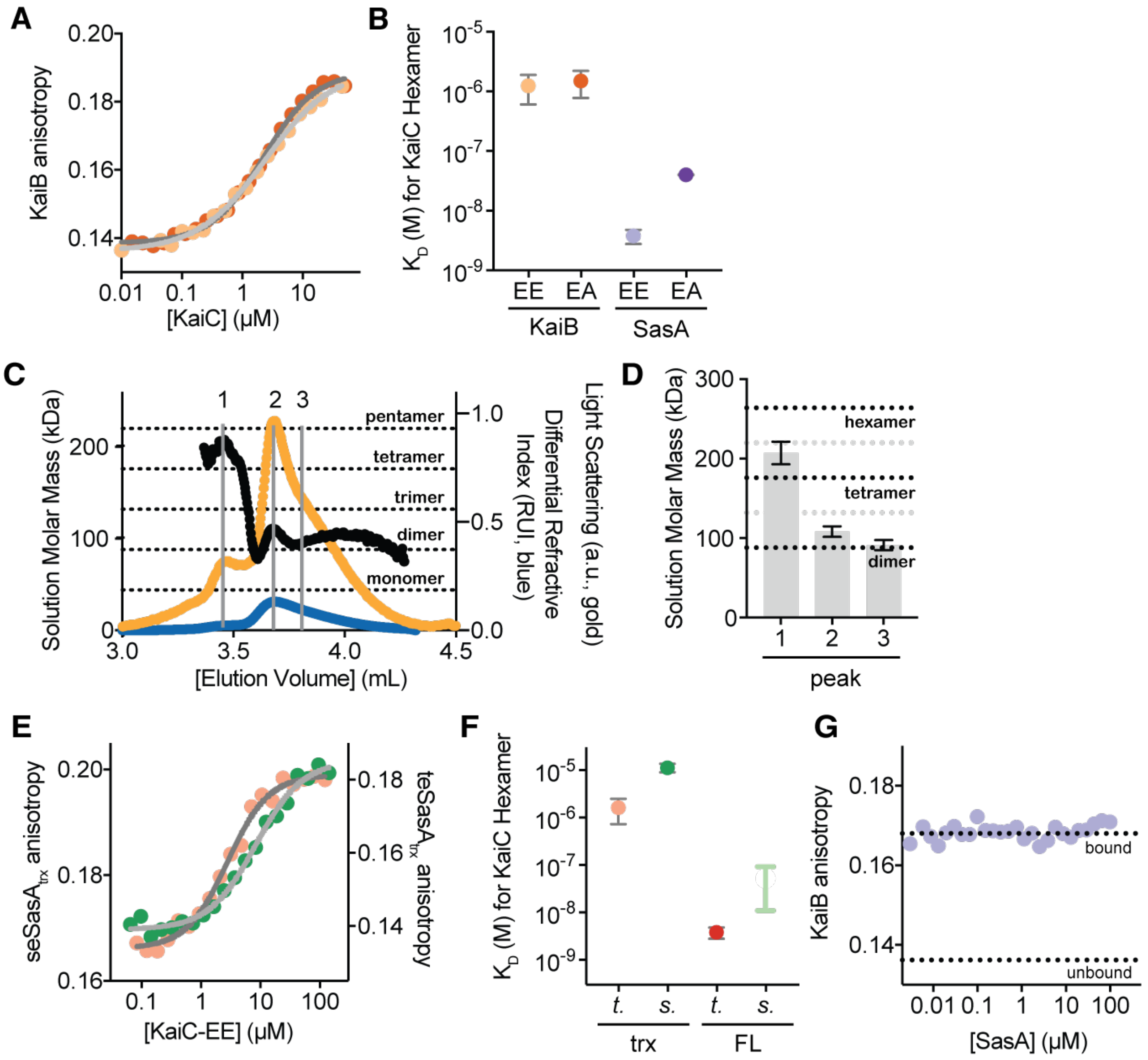

**Figure S2. Biochemical properties of the SasA-KaiC interaction.**

**A**) KaiB binds to KaiC-EE and KaiC-EA with similar affinity. Representative equilibrium binding assays of 50 nM fluorescently-labeled KaiB from *S. elongatus* in the presence of increasing concentrations of *S. elongatus* KaiC-EA (dark orange) and KaiC-EE (light orange). Data were fit to a Langmuir binding isotherm (EA, dark gray; EE, light gray). **B**) SasA binds KaiC more tightly than does KaiB, with a preference for the pS/pT phosphomimetic, KaiC-EE. Calculated equilibrium dissociation constants ($K_D$) from binding data in panel **A** (mean ± SD, n = 3; light and dark orange) are compared to affinities from Valencia et al. (26) (mean ± SD, $K_D$ measured by SPR; light and dark purple) for full-length SasA from *T. elongatus*. **C**) Full-length SasA exists primarily a dimer in solution. Traces from a representative size-exclusion chromatography coupled to multiangle light scattering (SEC-MALS) run for full-length SasA from *S. elongatus*. Differential refractive index (blue) is depicted with light scattering (gold) and absolute mass estimation (kDa, black line). **D**) Quantitative mass analysis of peaks indicated at the top of panel **C** (peaks 1-3) analyzed from triplicate SEC-MALS runs are consistent with a dimer-tetramer equilibrium (mean mass in kDa ± SD). Calculated masses for dimer, tetramer and hexamer are represented by black dotted lines. **E**) The $SasA_{trx}$ domain of *T. elongatus* binds with higher affinity than the *S. elongatus* $SasA_{trx}$ domain to the respective KaiC-EE variant from each organism. Equilibrium binding assays of fluorescently-labeled $SasA_{trx}$ from *S. elongatus* (dark green) or *T.* (pink) *elongatus*. Data were fit as in panel **A**. **F**) Full-length SasA (FL) binds much more tightly than the isolated $SasA_{trx}$ domain (trx), suggesting that it relies on avidity for efficient binding to KaiC hexamer. Calculated $K_D$ values extracted from data in panel **E** (mean ± SD, n = 3; pink or dark green) are compared to affinities from Valencia et al. (26) (mean ± SD, measured by SPR; red) for full-length SasA from *T. elongatus*. The affinity range of *S. elongatus* SasA calculated from the 2D titration data and our thermodynamic model (**Fig. 3F** and **S4**) is represented as light green bars. **G**) Full-length SasA does not display avidity and compete KaiB off of the KaiC-C1 monomer. Fluorescently-labeled KaiB was bound to 10 μM monomeric KaiC-CI domain and subsequently titrated with full-length SasA as indicated. Anisotropy values for bound and unbound KaiB (dashed lines) were determined from curve fits in **Fig. S4A**.

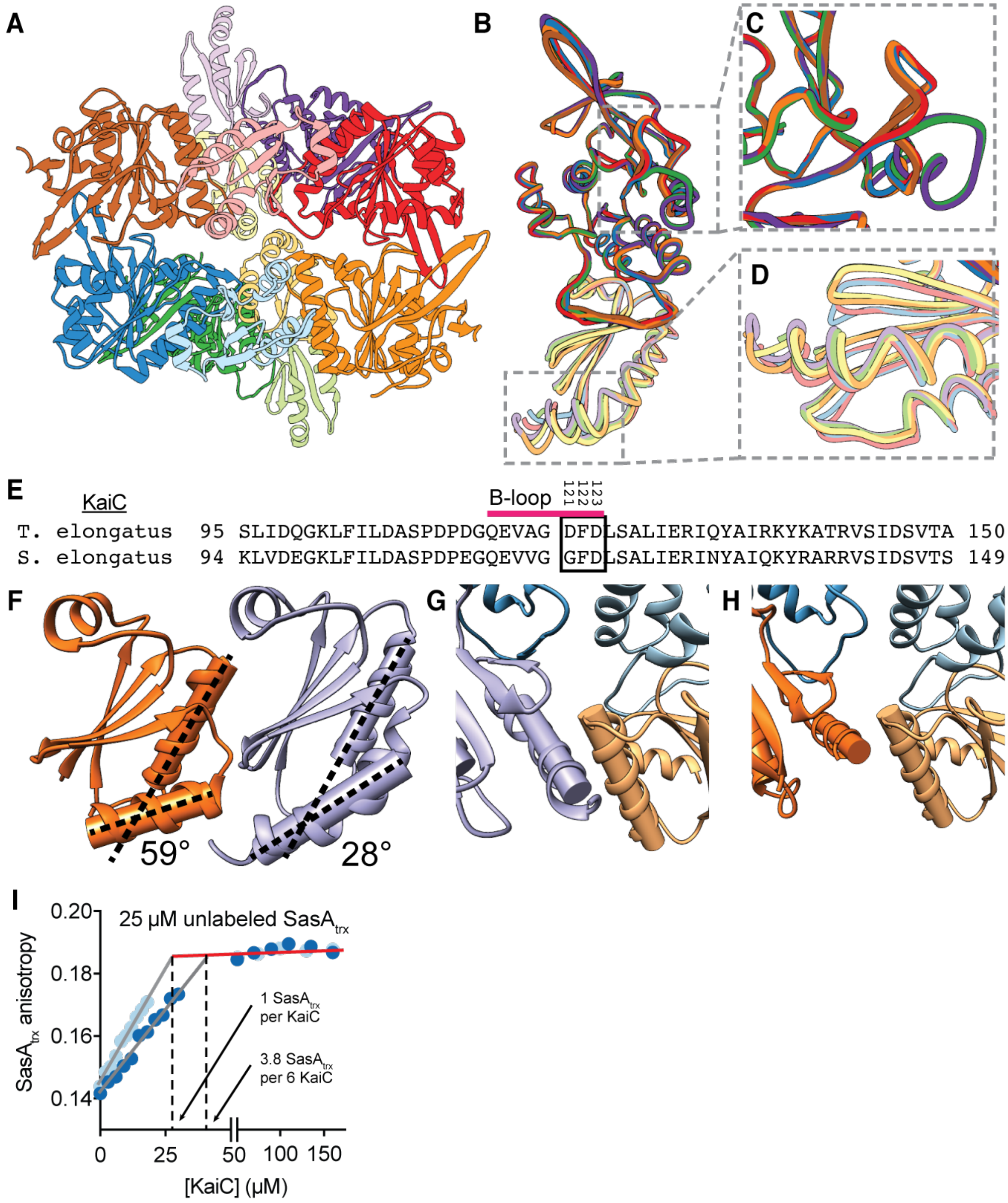

**Figure S3. Structural details of the KaiC CI-SasA$_{trx}$ complex.**

**A**) Asymmetric unit of the KaiC-C1-SasA$_{trx}$ crystal complex (PDB 6X61). Individual KaiC-CI-SasA$_{trx}$ pairs are depicted with SasA$_{trx}$ subunits in light hues and KaiC-CI subunits in dark hues. **B-D**) Modest structural heterogeneity was observed between the complexes of the asymmetric unit. Backbone overlays of the 6 KaiC CI-SasA$_{trx}$ complexes from the asymmetric unit. Small differences are highlighted by the orientation of a loop comprising residues R185-V201 on the CI domain (panel **C**) or by the orientation of the α3 helix of SasA$_{trx}$ (panel **D**), with consensus among the complexes of chains CD and KL. **E**) Multiple sequence alignment of the B-loops (pink) and surrounding sequence from KaiC of *S. elongatus* and *T. elongatus*. Black box, residues at the shared KaiB and SasA-binding interface that were subjected to mutagenesis **F**) SasA and fsKaiB diverge structurally at the α3 helix. The α1 and α3 helices of fsKaiB (PDB: 5JWO) and $SasA_{trx}$ (PDB: 6X61) are modeled as cylindrical axes with a radius of 1.8 Å, with the inter-axis crossing angle reported. **G-H**) Orientation of the α3 helix is likely to affect binding of KaiB at the adjacent KaiC protomer. Helical orientation of the $SasA_{trx}$ (purple, panel **G**) or fsKaiB (orange, panel **H**) domain relative to the α1 helix on the adjacent KaiB molecule at the CW interface (yellow). Helices are represented as helical axes as in panel **F**. **I**) Representative saturation binding titrations of the *T. elongatus* KaiC-CI monomer or KaiC-EE hexamer in the presence of 25 μM unlabeled $SasA_{trx}$. Stoichiometry was determined assuming 1:1 binding for $SasA_{trx}$ to the KaiC-CI monomer. Stoichiometry of the saturated KaiC-$SasA_{trx}$ complex was calculated at 3.8 ± 0.3 (mean ± SD, n = 3) molecules of SasA per KaiC hexamer, consistent with measurements from native mass spectrometry (27).

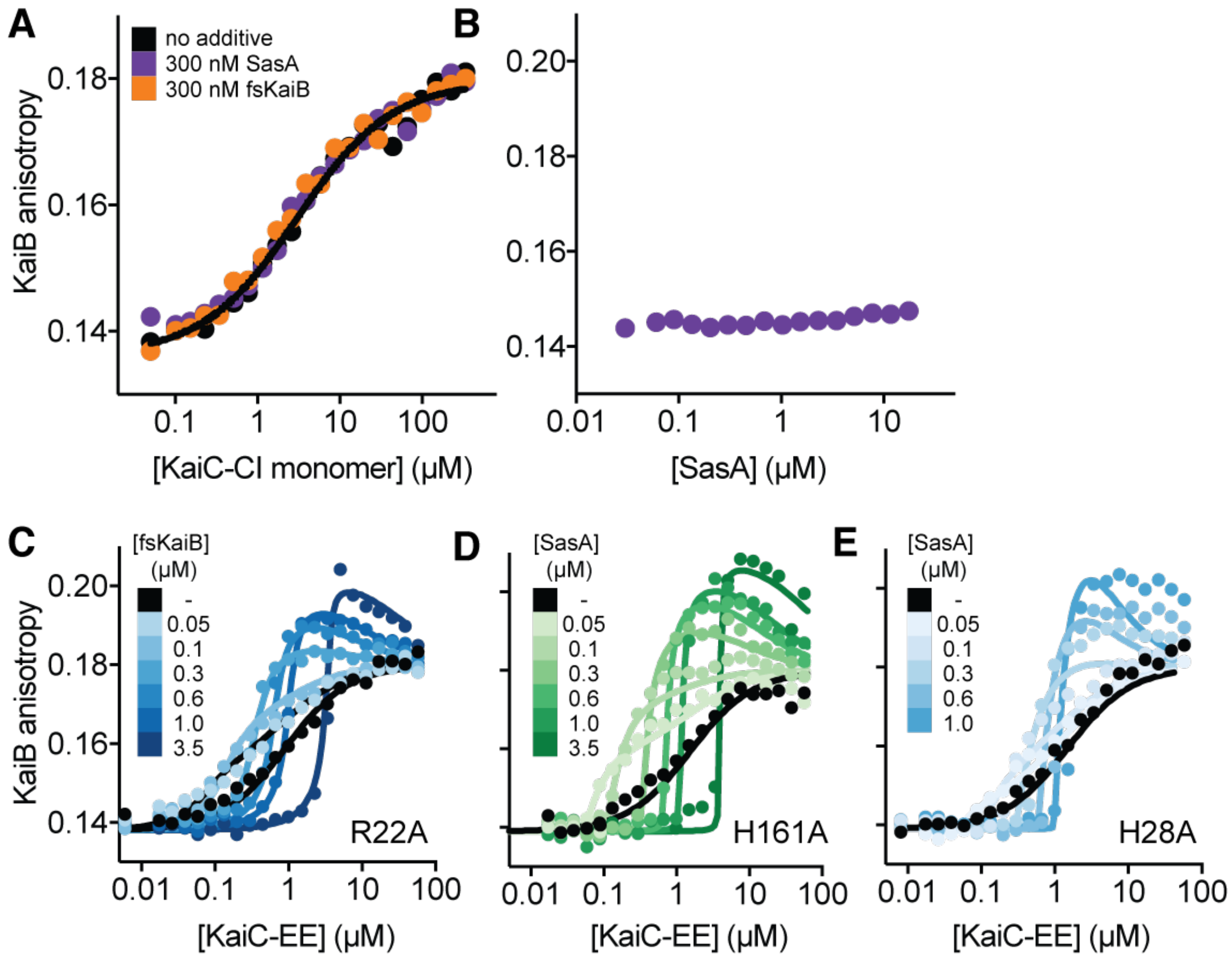

**Figure S4. Modeling and validating heterocooperative association of KaiB with KaiC.**

**A**) SasA and fsKaiB additives do not affect binding of 50 nM fluorescently-labeled KaiB to KaiC-CI monomer. Equilibrium binding titration of fluorescently-labeled KaiB with *S. elongatus* KaiC-CI monomer (black) in the presence of 300 nM fsKaiB (orange) or full-length SasA (purple). **B**) KaiC is required for interaction between SasA and KaiB. Equilibrium titration of fluorescently-labeled KaiB with full-length SasA (purple), showing that KaiB anisotropy is unaffected without KaiC present. **C-E**) 2D titration assays and associated fits from the thermodynamic model for fsKaiB-R22A (panel **C**), SasA-H161A (panel **D**), and SasA-H28A (panel **E**).

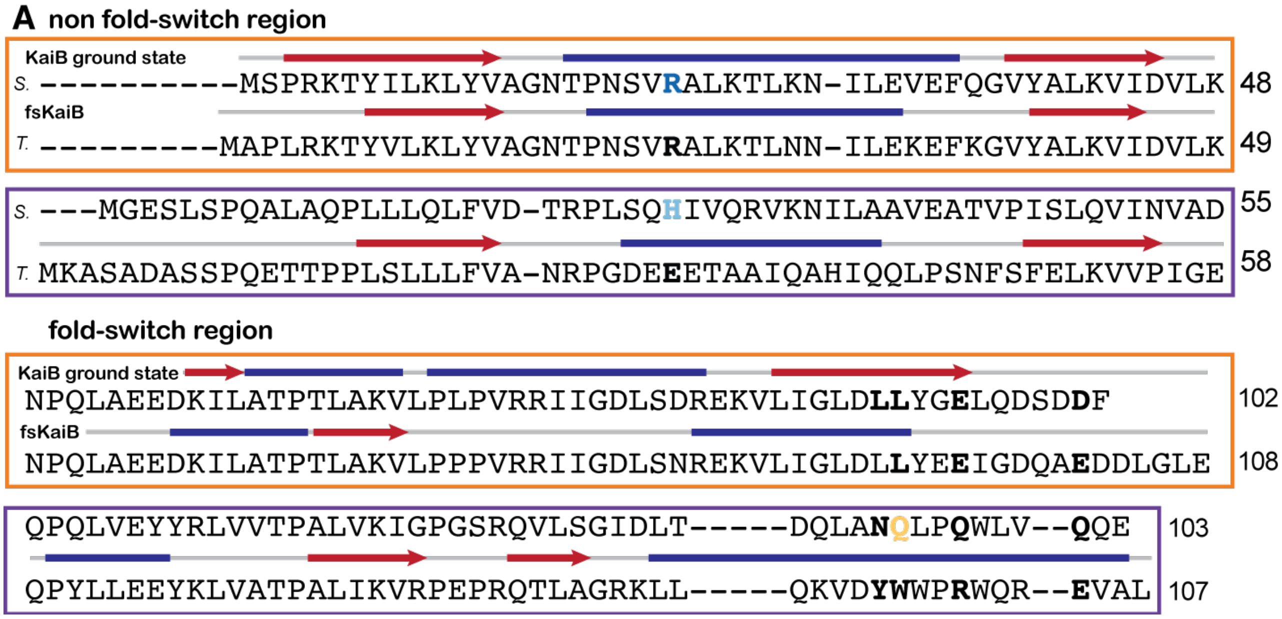

**Figure S5. Multiple sequence alignment of KaiB and SasA$_{trx}$ from *S.* and *T. elongatus*.**

**A**) Multiple sequence alignment of N-terminal (non-fold-switch) and C-terminal (fold-switch) halves of KaiB (orange box) and the structurally analogous SasA$_{trx}$ (purple box) from *S. elongatus* and *T. elongatus*. Red arrows (β-sheets) and blue lines (α-helices) indicate secondary structure from the KaiB tetramer (PDB: 2QKE), fsKaiB monomer (PDB: 5JWO) and SasA$_{trx}$ from the KaiC-CI-SasA$_{trx}$ structure (PDB: 6X61) reported here. Mutations tested in this study are depicted in bold.

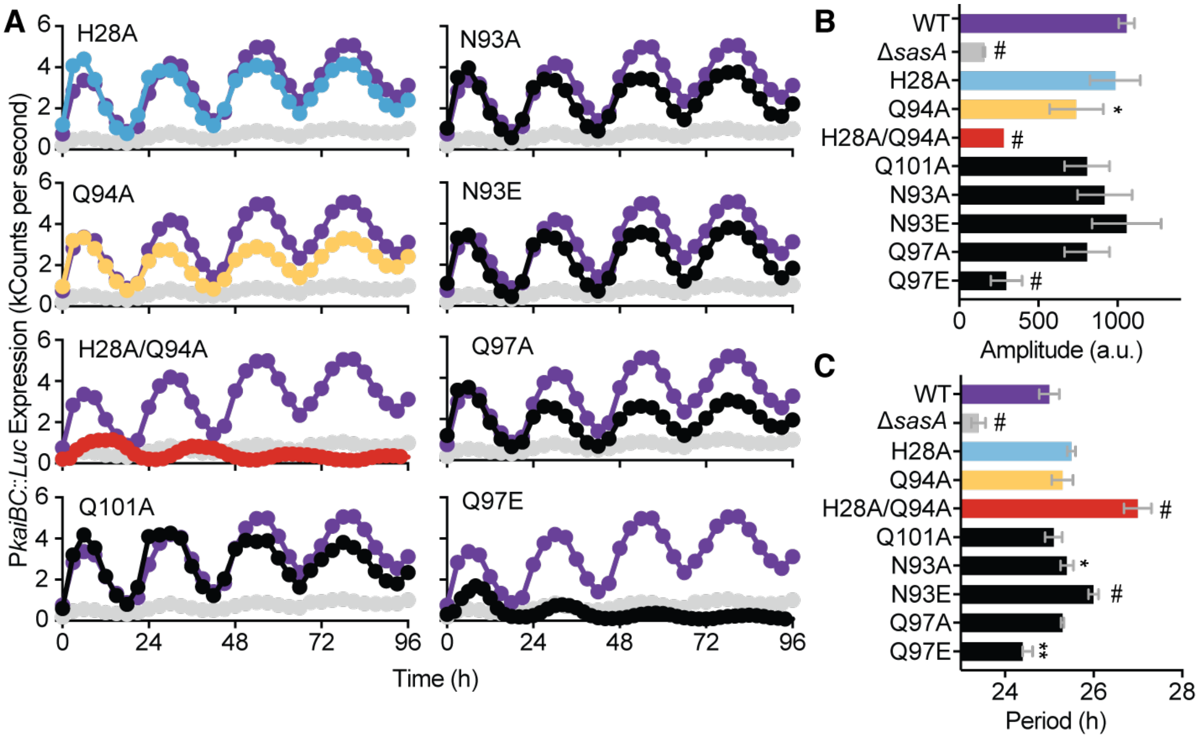

**Figure S6. Mutations at the SasA-KaiB cooperativity interface alter circadian rhythms in *S. elongatus*.**

**A**) Bioluminescence traces produced by *S. elongatus* strains based on expression of firefly luciferase driven by the clock-controlled *kaiBC* promoter. Wild type (WT) and Δ*sasA* strains are compared to markerless CRISPR/Cas12a-edited *sasA* mutants, with the relevant amino acid substitutions indicated. Graphs display the average of 6-12 wells, with standard deviation omitted for better visibility of multiple traces. WT (purple) and mutants, colored as in panel **B**. **B-C**) Analysis of raw bioluminescence data was performed using BioDare2 to calculate the signal amplitude (in arbitrary units, a.u.) (panel **B**) and period length (panel **C**), with the average and standard deviation reported for 6-12 replicates. One-way ANOVA was performed to identify significant changes (*, $P < 0.05$; **, $P < 0.01$; #, $P < 0.0001$) in amplitude and period length relative to wild type.

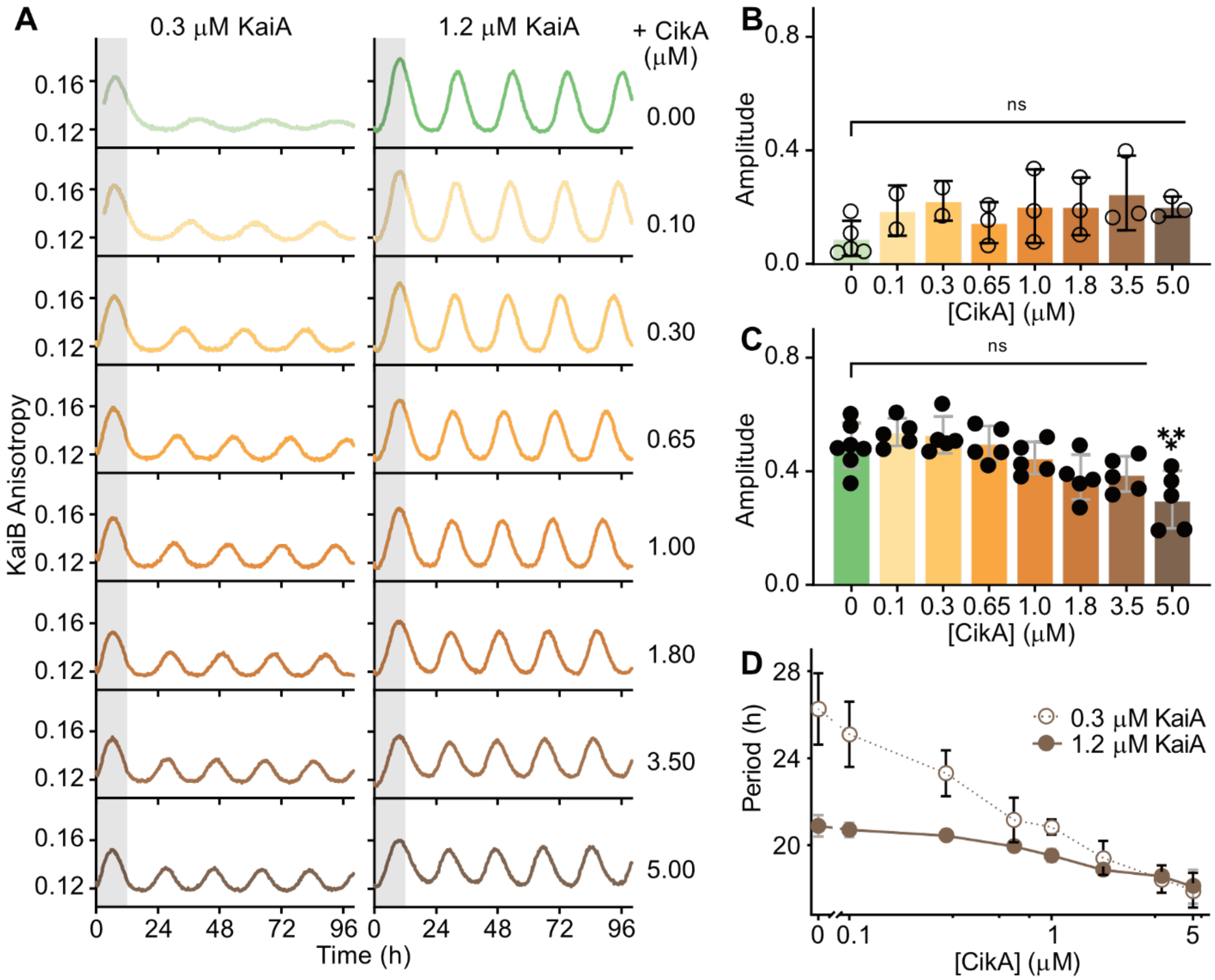

**Figure S7. Enhancement of oscillator robustness by CikA under limiting concentrations of KaiA.**

**A**) The FP-PTO under conditions of 3.5 μM KaiC, 3.5 μM KaiB (including 50 nM fluorescently-labeled KaiB as a probe), and either 0.3 μM or 1.2 μM KaiA as indicated with titrations of CikA from 0.1 – 5.0 μM. Representative assay from n ≥ 2 shown; the first 12-h period after release into constant conditions is marked in gray. **B** and **C**) Amplitude for FP-PTO assays prepared under different KaiA concentrations of (panel **B**) 0.3 μM (open circles) or (panel **C**) 1.2 μM (closed circles) KaiA in the absence (green) or presence of CikA (light yellow to brown). ANOVA was used to compare amplitudes of the FP-PTO under the two KaiA concentrations in the absence of CikA versus the indicated concentrations of CikA: ns, not significant; *, $P < 0.05$; **, $P < 0.01$; ***, $P < 0.001$; #, $P < 0.0001$. **D**) Period values for FP-PTO assays prepared with 0.3 μM (open circles) or 1.2 μM (closed circles) KaiA in the absence and presence of CikA (brown). Period and amplitude data are representative of two or more independent experiments, presented as the mean ± SD.

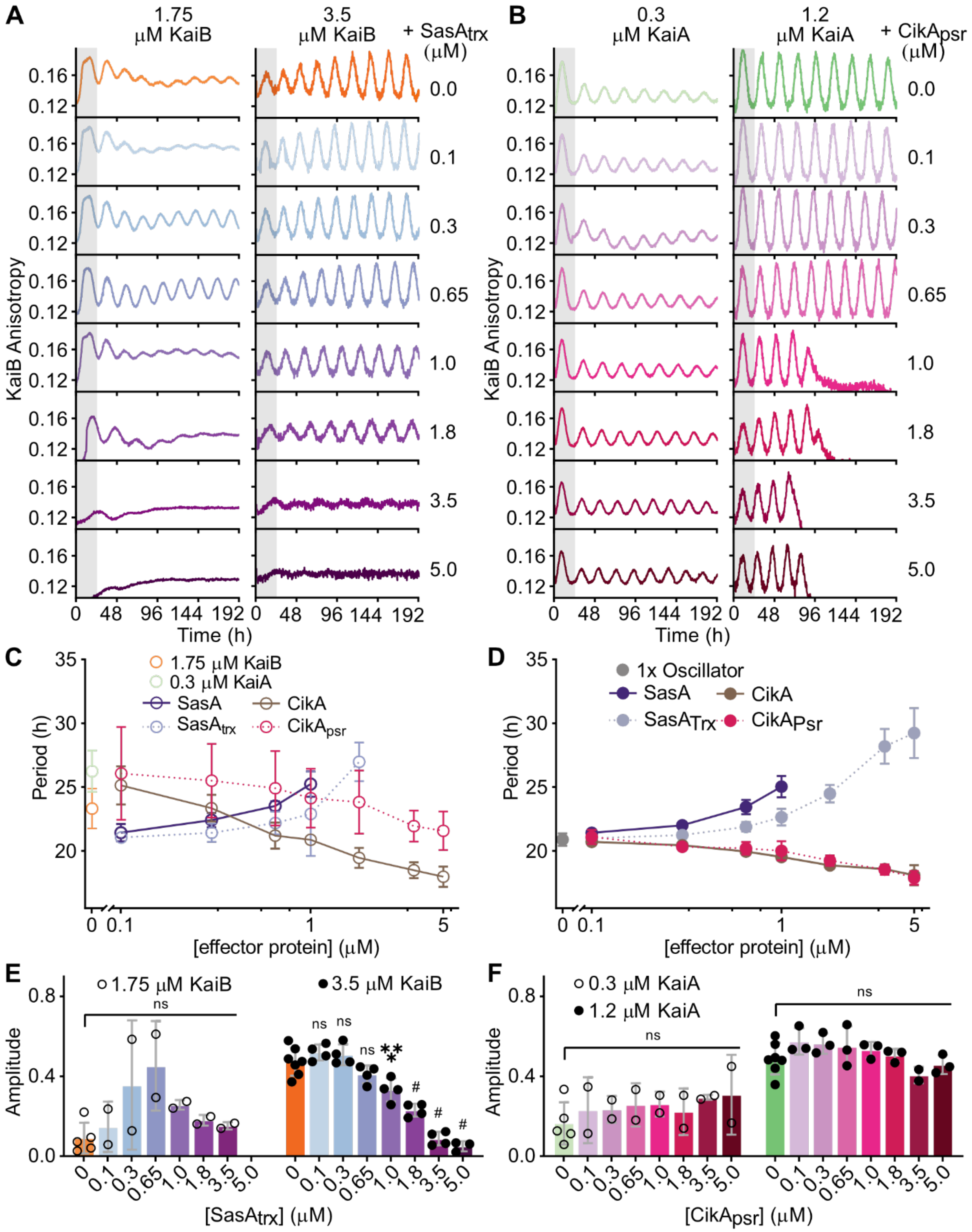

**Figure S8. Domain truncation studies on SasA and CikA oscillator effects.**

**A-B)** The FP-PTO under conditions of 3.5 μM KaiC, 50 nM fluorescently-labeled KaiB as a probe, and either (panel **A**) standard 1.2 μM KaiA with 1.75 μM or 3.5 μM KaiB, or (panel **B**) standard 3.5 μM KaiB with 0.3 μM or 1.2 μM KaiA as indicated with titrations of truncated domain (panel **A**) $SasA_{trx}$ or (panel **B**) $CikA_{psr}$ from 0.1 – 5.0 μM. Representative assay from n ≥ 2 shown; the first 24-h period after release into constant conditions is marked in gray. **C-D**) Period values for FP-PTO assays prepared with deficient (panel **C**) 1.75 μM KaiB and (panel **D**) 0.3 μM KaiA (open circles) or standard oscillator concentrations (panel **C**) 3.5 μM KaiB and (panel **D**) 1.2 μM KaiA (closed circles) in the absence and presence of full-length (solid connecting lines) and truncated additive proteins (dotted connecting lines). Period and amplitude data are representative of one or more independent experiments with duplicate samples, presented as the mean ± SD. **E-F)** Amplitude for FP-PTO assays prepared under different core KaiB and KaiA concentrations of (panel **E**) deficient (open circles) or (panel **F**) standard (closed circles) in the absence or presence of truncated additive proteins. Data are shown as mean ± SD (n ≥ 2 with duplicate samples). Analysis of variance (ANOVA) was used to compare amplitudes of the FP-PTO under the two KaiB and KaiA concentrations in the absence of truncated domains $SasA_{trx}$ and $CikA_{psr}$, respectively, versus the indicated concentrations of those additive proteins: ns, not significant; *, $P < 0.05$; **, $P < 0.01$; ***, $P < 0.001$; #, $P < 0.0001$.

**Table S1. Constructs and shorthand names used in this study.**

| Shorthand name | organism | Protein full name |
|---|---|---|
| KaiC | *S. elongatus* | FLAG-seKaiC-1-518 |
| KaiC-EE | | FLAG-seKaiC-1-518-S431E-T432E |
| | *T. elongatus* | FLAG-teKaiC-1-517-S431E-T432E |
| KaiC-EA | *S. elongatus* | FLAG-seKaiC-1-518-S431E-T432A |
| KaiC-EE-D121A | *T. elongatus* | FLAG-teKaiC-1-517-D121A-S431E-T432E |
| KaiC-EE-F122A | | FLAG-teKaiC-1-517-F122A-S431E-T432E |
| KaiC-EE-D123A | | FLAG-teKaiC-1-517-D123A-S431E-T432E |
| KaiC-CI monomer | | FLAG-teKaiC-17-247-R41A-K173A |
| | | FLAG-seKaiC-16-246-R40A-K172A |
| KaiB | *S. elongatus* | seKaiB-1-102-FLAG |
| KaiB-K25C-6IAF | | sekaiB-1-102-K25C-FLAG-fluorescein |
| fsKaiB | | seKaiB-1-99-Y7A-I87A-Y93A-FLAG |
| fsKaiB-R22A | | seKaiB-1-99-Y7A-R22A-I87A-Y93A-FLAG |
| KaiA | | seKaiA-1-284 |
| SasA | | seSasA-1-387-FLAG |
| SasA-H161A | | seSasA-1-387-H161A-FLAG |
| SasA-H28A | | seSasA-1-387-H28A-FLAG |
| SasA-Q94A | | seSasA-1-387-Q94A-FLAG |
| SasA-H28A-Q94A | | seSasA-1-387-H28A-Q94A-FLAG |
| SasA$_{trx}$ | | FLAG-seSasA-13-103-P13A-FLAG |
| | *T. elongatus* | FLAG-teSasA-16-107-P16A-FLAG |
| SasA$_{trx}$-Q31C* | *S. elongatus* | FLAG-seSasA-13-103-P13A-Q31C-FLAG |
| SasA$_{trx}$-A35C* | *T. elongatus* | FLAG-teSasA-16-107-P16A-A35C-FLAG |
| CikA | *S. elongatus* | FLAG-seCikA-1-754 |
| CikA$_{psr}$ | | seCikA-S605-606-745-C644S-C686S-C742S |
| CikA$_{psr}$-S727C* | | seCikA-S605-606-745-C644S-C686S-C742S-S727C |

* single cysteine residue for labeling with 6-iodoacetamido-fluorescein

**Table S2. Specific reaction conditions with designated protein constructs.**

| Experiment Type | Protein Construct Shorthand Name / Final Concentration (µM if not indicated) | Experimental Condition |
|---|---|---|
| KaiC Phosphorylation Assay (Fig. 1B, S1) | 1) KaiC / 3.5<br>2) KaiB / 3.5<br>3) KaiA / 1.2 | • Volume: 1.8 mL<br>• Buffer: 20 mM Tris, 150 mM NaCl, 5 mM $MgCl_2$, 1 mM ATP, 0.5 mM EDTA, pH 8.0<br>• Temperature: 30 °C |
| FP-PTO Assay tracking fluorescently labeled KaiB (Fig. 1B, S1) | 1) KaiB-K25C-6IAF / 0.05<br>2) KaiC / 3.5<br>3) KaiB / 3.45<br>4) KaiA / 1.2 | • Volume: 80 µL<br>• Buffer: 20 mM Tris, 150 mM NaCl, 5 mM $MgCl_2$, 1 mM ATP, 0.5 mM EDTA, pH 8.0<br>• Temperature: 30 °C |
| FP-PTO Assay tracking fluorescently labeled $SasA_{trx}$ (Fig. 1B, S1) | 1) $SasA_{trx}$-Q31C-6IAF / 0.05<br>2) KaiC / 3.5<br>3) KaiB / 3.5<br>4) KaiA / 1.2 | • Volume: 80 µL<br>• Buffer: 20 mM Tris, 150 mM NaCl, 5 mM $MgCl_2$, 1 mM ATP, 0.5 mM EDTA, pH 8.0<br>• Temperature: 30 °C |
| KaiA Titration FP-PTO Assay (Fig. 4A) | 1) KaiB-K25C-6IAF / 0.05<br>2) KaiC / 3.5<br>3) KaiB / 3.45<br>4) KaiA / varied (0.3, 0.6, 1.2, 2.4, 3.6, 4.8, and 6.0) | • Volume: 80 µL<br>• Buffer: 20 mM Tris, 150 mM NaCl, 5 mM $MgCl_2$, 1 mM ATP, 0.5 mM EDTA, pH 8.0<br>• Temperature: 30 °C |
| KaiB Titration FP-PTO Assay (Fig. 4B) | 1) KaiB-K25C-6IAF / 0.05<br>2) KaiC / 3.5<br>3) KaiB / varied (0.65, 0.825, 1.70, 3.45, 6.95, 10.45, and 17.45)<br>4) KaiA / 1.2 | • Volume: 80 µL<br>• Buffer: 20 mM Tris, 150 mM NaCl, 5 mM $MgCl_2$, 1 mM ATP, 0.5 mM EDTA, pH 8.0<br>• Temperature: 30 °C |
| SasA Titration FP-PTO Assay under different KaiB concentrations (Fig. 5) | 1) KaiB-K25C-6IAF / 0.05<br>2) KaiC / 3.5<br>3) KaiB / fixed at 0.825, 1.70, and 3.45<br>4) KaiA / 1.2<br>5) SasA / varied (0.0, 0.1, 0.65, 1.0, 1.8, 3.5, and 5.0) | • Volume: 80 µL<br>• Buffer: 20 mM Tris, 150 mM NaCl, 5 mM $MgCl_2$, 1 mM ATP, 0.5 mM EDTA, pH 8.0<br>• Temperature: 30 °C |
| CikA Titration FP-PTO Assay under different KaiA concentrations (S6) | 1) KaiB-K25C-6IAF / 0.05<br>2) KaiC / 3.5<br>3) KaiB / 3.45<br>4) KaiA / fixed at 0.3 and 1.2<br>5) CikA / varied (0.0, 0.1, 0.65, 1.0, 1.8, 3.5, and 5.0) | • Volume: 80 µL<br>• Buffer: 20 mM Tris, 150 mM NaCl, 5 mM $MgCl_2$, 1 mM ATP, 0.5 mM EDTA, pH 8.0<br>• Temperature: 30 °C |
| $SasA_{trx}$ Titration FP-PTO Assay under different KaiB concentrations (S7A) | 1) KaiB-K25C-6IAF / 0.05<br>2) KaiC / 3.5<br>3) KaiB / fixed at 1.70 and 3.45<br>4) KaiA / 1.2<br>5) $SasA_{trx}$ / varied (0.0, 0.1, 0.65, 1.0, 1.8, 3.5, and 5.0) | • Volume: 80 µL<br>• Buffer: 20 mM Tris, 150 mM NaCl, 5 mM $MgCl_2$, 1 mM ATP, 0.5 mM EDTA, pH 8.0<br>• Temperature: 30 °C |
| $CikA_{psr}$ Titration FP-PTO Assay under different KaiA concentrations (S7B) | 1) KaiB-K25C-6IAF / 0.05<br>2) KaiC / 3.5<br>3) KaiB / 3.45<br>4) KaiA / fixed at 0.3 and 1.2<br>5) $CikA_{psr}$ / varied (0.0, 0.1, 0.65, 1.0, 1.8, 3.5, and 5.0) | • Volume: 80 µL<br>• Buffer: 20 mM Tris, 150 mM NaCl, 5 mM $MgCl_2$, 1 mM ATP, 0.5 mM EDTA, pH 8.0<br>• Temperature: 30 °C |

**Table S3. Refinement statistics for structure determination of KaiC-CI-$SasA_{trx}$ complex.**

| **Data collection** | |
|---|---|
| Space group | $P2_1$ |
| Cell dimensions | |
| *a*, *b*, *c* (Å) | 107.6, 121.58, 133.59 |
| α, β, γ (°) | 90.0, 108.78, 90.0 |
| Resolution (Å) | 49.05-3.2 (3.30-3.20) * |
| No. of total reflections | 153257 (13790) |
| No. Unique Reflections | 53267 (4637) |
| $R_{merge}$ | 16.5 (75.7) |
| $R_{pim}$ | 13.0 (57.5) |
| I/σI | 5.5 (1.5) |
| Completeness (%) | 98.8 (99.3) |
| $CC_{1/2}$ | 0.95 (0.58) |
| Wilson B-factor | 58.3 |
| Redundancy | 2.9 (3.0) |
| **Refinement** | |
| Resolution (Å) | 3.20 |
| $R_{\mathrm{work}}$/ $R_{\mathrm{free}}$ | 22.2/26.6 |
| No. atoms | |
| Protein | 15469 |
| Ligand/ion | 30 |
| B-factors | |
| Protein | 61.3 |
| Ligand/ion | 46.1 |
| R.m.s deviations | |
| Bond lengths (Å) | 0.003 |
| Bond angles (°) | 0.78 |
| Residues in favored regions (%) | 93.1 |
| Residues in outlier regions (%) | 0.6 |

*Highest resolution shell is shown in parenthesis.
One crystal was used for data collection.
Data were collected at a wavelength of 1.0 Å.

### Table S4. Period analysis, ordinary one-way ANOVA for FP-PTO assays comparing different oscillator conditions.

| Protein-concentration (μM) comparison with / altered protein-concentration | Mean Diff. | 95.00% CI of diff. | Significant? | Summary | Adjusted *P* Value |
|---|---|---|---|---|---|
| KaiB-3.5 / KaiB-1.75 | -1.666 | -3.238 to -0.09300 | Yes | * | 0.0354 |
| KaiB-3.5 / KaiB-7.0 | 0.5423 | -1.394 to 2.479 | No | ns | 0.8847 |
| KaiB-3.5 / KaiB-10.5 | 0.5118 | -1.561 to 2.584 | No | ns | 0.922 |
| KaiB-3.5 / KaiB-17.5 | 1.244 | -0.6920 to 3.181 | No | ns | 0.3047 |
| SasA-0.0 / SasA-0.1 | -0.498 | -1.678 to 0.6824 | No | ns | 0.622 |
| SasA-0.0 / SasA-0.3 | -1.086 | -2.266 to 0.09437 | No | ns | 0.0758 |
| SasA-0.0 / SasA-0.65 | -2.52 | -3.700 to -1.340 | Yes | *** | 0.0001 |
| SasA-0.0 / SasA-1.0 | -4.118 | -5.298 to -2.938 | Yes | # | <0.0001 |
| SasA-0.0 / SasA-1.8 | -12.76 | -14.49 to -11.03 | Yes | # | <0.0001 |
| KaiB-1.75, SasA-0.0 / SasA-0.1 | 1.953 | 0.5292 to 3.377 | Yes | ** | 0.0048 |
| KaiB-1.75, SasA-0.0 / SasA-0.3 | 0.9688 | -0.5609 to 2.498 | No | ns | 0.3415 |
| KaiB-1.75, SasA-0.0 / SasA-0.65 | -0.1362 | -1.666 to 1.393 | No | ns | 0.9997 |
| KaiB-1.75, SasA-0.0 / SasA-1.0 | -1.861 | -3.285 to -0.4367 | Yes | ** | 0.0074 |
| KaiB-1.75, SasA-0.0 / SasA-1.8 | -8.009 | -10.66 to -5.359 | Yes | # | <0.0001 |
| $SasA_{trx}$-0.0 / $SasA_{trx}$-0.1 | -0.1257 | -1.638 to 1.387 | No | ns | 0.9997 |
| $SasA_{trx}$-0.0 / $SasA_{trx}$-0.3 | -0.3682 | -1.881 to 1.144 | No | ns | 0.9812 |
| $SasA_{trx}$-0.0 / $SasA_{trx}$-0.65 | -1.008 | -2.521 to 0.5041 | No | ns | 0.3257 |
| $SasA_{trx}$-0.0 / $SasA_{trx}$-1.0 | -1.771 | -3.283 to -0.2584 | Yes | * | 0.016 |
| $SasA_{trx}$-0.0 / $SasA_{trx}$-1.8 | -3.611 | -5.123 to -2.098 | Yes | # | <0.0001 |
| $SasA_{trx}$-0.0 / $SasA_{trx}$-3.5 | -7.318 | -8.831 to -5.806 | Yes | # | <0.0001 |
| $SasA_{trx}$-0.0 / $SasA_{trx}$-5.0 | -8.366 | -10.03 to -6.701 | Yes | # | <0.0001 |
| KaiA-1.2 / KaiA-0.3 | -5.433 | -6.896 to -3.969 | Yes | # | <0.0001 |
| KaiA-1.2 / KaiA-0.6 | -1.293 | -2.833 to 0.2473 | No | ns | 0.1285 |
| KaiA-1.2 / KaiA-2.4 | 0.5891 | -0.9510 to 2.129 | No | ns | 0.8174 |
| KaiA-1.2 / KaiA-3.6 | 0.9321 | -0.7165 to 2.581 | No | ns | 0.4766 |
| KaiA-1.2 / KaiA-4.8 | -1.593 | -3.702 to 0.5161 | No | ns | 0.2017 |
| KaiA-1.2 / KaiA-6.0 | -4.173 | -6.985 to -1.361 | Yes | ** | 0.002 |
| CikA-0.0 / CikA-0.1 | -0.04685 | -0.1630 to 0.06934 | No | ns | 0.811 |
| CikA-0.0 / CikA-0.3 | -0.03718 | -0.1534 to 0.07901 | No | ns | 0.9261 |
| CikA-0.0 / CikA-0.65 | -0.00668 | -0.1229 to 0.1095 | No | ns | 0.9998 |
| CikA-0.0 / CikA-1.0 | 0.04403 | -0.07215 to 0.1602 | No | ns | 0.85 |
| CikA-0.0 / CikA-1.8 | 0.1113 | -0.004878 to 0.2275 | No | ns | 0.0654 |
| CikA-0.0 / CikA-3.5 | 0.1006 | -0.01555 to 0.2168 | No | ns | 0.1145 |
| CikA-0.0 / CikA-5.0 | 0.1906 | 0.07439 to 0.3068 | Yes | *** | 0.0004 |
| KaiA-0.3, CikA-0.0 / CikA-0.1 | 0.75 | -2.315 to 3.815 | No | ns | 0.9499 |
| KaiA-0.3, CikA-0.0 / CikA-0.3 | 2.545 | -0.5204 to 5.610 | No | ns | 0.1201 |
| KaiA-0.3, CikA-0.0 / CikA-0.65 | 4.693 | 1.895 to 7.492 | Yes | ** | 0.0014 |
| KaiA-0.3, CikA-0.0 / CikA-1.0 | 5.02 | 2.222 to 7.818 | Yes | *** | 0.0007 |

| | | | | | |
|---|---|---|---|---|---|
| KaiA-0.3, CikA-0.0 / CikA-1.8 | 6.44 | 3.642 to 9.238 | Yes | # | <0.0001 |
| KaiA-0.3, CikA-0.0 / CikA-3.5 | 7.407 | 4.608 to 10.21 | Yes | # | <0.0001 |
| KaiA-0.3, CikA-0.0 / CikA-5.0 | 7.923 | 5.125 to 10.72 | Yes | # | <0.0001 |
| $CikA_{psr}$-0.0 / $CikA_{psr}$-0.1 | 3.01 | 2.132 to 3.889 | Yes | # | <0.0001 |
| $CikA_{psr}$-0.0 / $CikA_{psr}$-0.3 | 2.361 | 1.386 to 3.336 | Yes | # | <0.0001 |
| $CikA_{psr}$-0.0 / $CikA_{psr}$-0.65 | 1.658 | 0.7792 to 2.536 | Yes | # | <0.0001 |
| $CikA_{psr}$-0.0 / $CikA_{psr}$-1.0 | 0.8853 | 0.006697 to 1.764 | Yes | * | 0.0476 |
| $CikA_{psr}$-0.0 / $CikA_{psr}$-1.8 | 0.6953 | -0.1833 to 1.574 | No | ns | 0.1771 |
| $CikA_{psr}$-0.0 / $CikA_{psr}$-3.5 | 0.5711 | -0.4036 to 1.546 | No | ns | 0.4758 |
| $CikA_{psr}$-0.0 / $CikA_{psr}$-5.0 | -0.1997 | -1.078 to 0.6789 | No | ns | 0.9893 |
| KaiB-1.75, $SasA_{trx}$-0.0 / $SasA_{trx}$-0.1 | 2.256 | -1.984 to 6.496 | No | ns | 0.4607 |
| KaiB-1.75, $SasA_{trx}$-0.0 / $SasA_{trx}$-0.3 | 1.906 | -2.334 to 6.146 | No | ns | 0.6179 |
| KaiB-1.75, $SasA_{trx}$-0.0 / $SasA_{trx}$-0.65 | 1.131 | -3.109 to 5.371 | No | ns | 0.9285 |
| KaiB-1.75, $SasA_{trx}$-0.0 / $SasA_{trx}$-1.0 | 0.416 | -3.824 to 4.656 | No | ns | 0.9996 |
| KaiB-1.75, $SasA_{trx}$-0.0 / $SasA_{trx}$-1.8 | -3.684 | -7.924 to 0.5561 | No | ns | 0.0974 |
| KaiB-1.75, $SasA_{trx}$-0.0 / $SasA_{trx}$-3.5 | -11.24 | -15.48 to -6.999 | Yes | # | <0.0001 |
| KaiA-0.3, $CikA_{psr}$-0.0 / $CikA_{psr}$-0.1 | -0.065 | -6.407 to 6.277 | No | ns | >0.9999 |
| KaiA-0.3, $CikA_{psr}$-0.0 / $CikA_{psr}$-0.3 | 0.505 | -5.837 to 6.847 | No | ns | 0.9997 |
| KaiA-0.3, $CikA_{psr}$-0.0 / $CikA_{psr}$-0.65 | 1.13 | -5.212 to 7.472 | No | ns | 0.9937 |
| KaiA-0.3, $CikA_{psr}$-0.0 / $CikA_{psr}$-1.0 | 1.905 | -4.437 to 8.247 | No | ns | 0.9136 |
| KaiA-0.3, $CikA_{psr}$-0.0 / $CikA_{psr}$-1.8 | 2.21 | -4.132 to 8.552 | No | ns | 0.845 |
| KaiA-0.3, $CikA_{psr}$-0.0 / $CikA_{psr}$-3.5 | 4.11 | -2.232 to 10.45 | No | ns | 0.3006 |
| KaiA-0.3, $CikA_{psr}$-0.0 / $CikA_{psr}$-5.0 | 4.47 | -1.872 to 10.81 | No | ns | 0.2302 |

Ordinary one-way ANOVA performed with Dunnett's multiple comparisons test was used to compare period values of the FP-PTO under specified conditions: ns, not significant; *, $P < 0.05$; **, $P < 0.01$; ***, $P < 0.001$; #, $P < 0.0001$. If not specified, FP-PTO assays were run with 1.2 μM KaiA, 3.45 μM KaiB, 3.5 μM KaiC, and 0.05 μM fluorescently-labeled KaiB probe.

**Table S5. Amplitude analysis, ordinary one-way ANOVA for FP-PTO assays comparing different oscillator conditions.**

| Protein-concentration (μM) comparison with / altered protein-concentration | Mean Diff. | 95.00% CI of diff. | Significant? | Summary | Adjusted *P* Value |
|---|---|---|---|---|---|
| KaiB-3.5 / KaiB-1.75 | 0.08743 | -0.02121 to 0.1961 | No | ns | 0.1437 |
| KaiB-3.5 / KaiB-7.0 | 0.04213 | -0.07417 to 0.1584 | No | ns | 0.7636 |
| KaiB-3.5 / KaiB-10.5 | 0.04656 | -0.06209 to 0.1552 | No | ns | 0.6484 |
| KaiB-3.5 / KaiB-17.5 | 0.434 | 0.3457 to 0.5222 | Yes | # | <0.0001 |
| SasA-0.0 / SasA-0.1 | -0.03844 | -0.1930 to 0.1162 | No | ns | 0.9098 |
| SasA-0.0 / SasA-0.3 | 0.02229 | -0.1323 to 0.1769 | No | ns | 0.989 |
| SasA-0.0 / SasA-0.65 | 0.1149 | -0.03974 to 0.2694 | No | ns | 0.1799 |
| SasA-0.0 / SasA-1.0 | 0.2204 | 0.06584 to 0.3750 | Yes | ** | 0.0047 |
| SasA-0.0 / SasA-1.8 | 0.4423 | 0.2160 to 0.6686 | Yes | *** | 0.0003 |
| KaiB-1.75, SasA-0.0 / SasA-0.1 | -0.4206 | -0.6723 to -0.1688 | Yes | *** | 0.0007 |
| KaiB-1.75, SasA-0.0 / SasA-0.3 | -0.5336 | -0.8040 to -0.2632 | Yes | # | <0.0001 |
| KaiB-1.75, SasA-0.0 / SasA-0.65 | -0.524 | -0.7944 to -0.2535 | Yes | *** | 0.0001 |
| KaiB-1.75, SasA-0.0 / SasA-1.0 | -0.3502 | -0.6019 to -0.09843 | Yes | ** | 0.0043 |
| KaiB-1.75, SasA-0.0 / SasA-1.8 | 0.007592 | -0.4608 to 0.4759 | No | ns | >0.9999 |
| $SasA_{trx}$-0.0 / $SasA_{trx}$-0.1 | -0.04557 | -0.1360 to 0.04490 | No | ns | 0.6219 |
| $SasA_{trx}$-0.0 / $SasA_{trx}$-0.3 | -0.03403 | -0.1245 to 0.05644 | No | ns | 0.8555 |
| $SasA_{trx}$-0.0 / $SasA_{trx}$-0.65 | 0.06425 | -0.02622 to 0.1547 | No | ns | 0.2642 |
| $SasA_{trx}$-0.0 / $SasA_{trx}$-1.0 | 0.1418 | 0.05135 to 0.2323 | Yes | *** | 0.0009 |
| $SasA_{trx}$-0.0 / $SasA_{trx}$-1.8 | 0.2437 | 0.1532 to 0.3341 | Yes | # | <0.0001 |
| $SasA_{trx}$-0.0 / $SasA_{trx}$-3.5 | 0.3859 | 0.2954 to 0.4763 | Yes | # | <0.0001 |
| $SasA_{trx}$-0.0 / $SasA_{trx}$-5.0 | 0.4243 | 0.3247 to 0.5239 | Yes | # | <0.0001 |
| KaiA-1.2 / KaiA-0.3 | 0.3983 | 0.1870 to 0.6096 | Yes | *** | 0.0001 |
| KaiA-1.2 / KaiA-0.6 | 0.1897 | -0.02157 to 0.4010 | No | ns | 0.0919 |
| KaiA-1.2 / KaiA-2.4 | 0.12 | -0.09133 to 0.3313 | No | ns | 0.4707 |
| KaiA-1.2 / KaiA-3.6 | 0.2376 | 0.01147 to 0.4638 | Yes | * | 0.0365 |
| KaiA-1.2 / KaiA-4.8 | 0.4218 | 0.1324 to 0.7111 | Yes | ** | 0.0025 |
| KaiA-1.2 / KaiA-6.0 | 0.4745 | 0.08869 to 0.8602 | Yes | * | 0.0115 |
| CikA-0.0 / CikA-0.1 | -0.04685 | -0.1630 to 0.06934 | No | ns | 0.811 |
| CikA-0.0 / CikA-0.3 | -0.03718 | -0.1534 to 0.07901 | No | ns | 0.9261 |
| CikA-0.0 / CikA-0.65 | -0.00668 | -0.1229 to 0.1095 | No | ns | 0.9998 |
| CikA-0.0 / CikA-1.0 | 0.04403 | -0.07215 to 0.1602 | No | ns | 0.85 |
| CikA-0.0 / CikA-1.8 | 0.1113 | -0.004878 to 0.2275 | No | ns | 0.0654 |
| CikA-0.0 / CikA-3.5 | 0.1006 | -0.01555 to 0.2168 | No | ns | 0.1145 |
| CikA-0.0 / CikA-5.0 | 0.1906 | 0.07439 to 0.3068 | Yes | *** | 0.0004 |
| KaiA-0.3, CikA-0.0 / CikA-0.1 | -0.09828 | -0.3229 to 0.1263 | No | ns | 0.7137 |
| KaiA-0.3, CikA-0.0 / CikA-0.3 | -0.1326 | -0.3571 to 0.09201 | No | ns | 0.4194 |
| KaiA-0.3, CikA-0.0 / CikA-0.65 | -0.05589 | -0.2519 to 0.1401 | No | ns | 0.9446 |
| KaiA-0.3, CikA-0.0 / CikA-1.0 | -0.1135 | -0.3095 to 0.08252 | No | ns | 0.4393 |

| | | | | | |
|---|---|---|---|---|---|
| KaiA-0.3, CikA-0.0 / CikA-1.8 | -0.1124 | -0.3085 to 0.08358 | No | ns | 0.449 |
| KaiA-0.3, CikA-0.0 / CikA-3.5 | -0.1599 | -0.3560 to 0.03607 | No | ns | 0.1412 |
| KaiA-0.3, CikA-0.0 / CikA-5.0 | -0.1116 | -0.3076 to 0.08441 | No | ns | 0.4568 |
| $CikA_{psr}$-0.0 / $CikA_{psr}$-0.1 | 0.01182 | -0.2952 to 0.3189 | No | ns | 0.9999 |
| $CikA_{psr}$-0.0 / $CikA_{psr}$-0.3 | -0.0595 | -0.3976 to 0.2786 | No | ns | 0.9971 |
| $CikA_{psr}$-0.0 / $CikA_{psr}$-0.65 | 0.02002 | -0.2870 to 0.3271 | No | ns | 0.9997 |
| $CikA_{psr}$-0.0 / $CikA_{psr}$-1.0 | 0.06736 | -0.2397 to 0.3744 | No | ns | 0.9899 |
| $CikA_{psr}$-0.0 / $CikA_{psr}$-1.8 | 0.08699 | -0.2201 to 0.3941 | No | ns | 0.9588 |
| $CikA_{psr}$-0.0 / $CikA_{psr}$-3.5 | 0.196 | -0.1420 to 0.5341 | No | ns | 0.4727 |
| $CikA_{psr}$-0.0 / $CikA_{psr}$-5.0 | 0.1054 | -0.2016 to 0.4125 | No | ns | 0.9001 |
| KaiB-1.75, $SasA_{trx}$-0.0 / $SasA_{trx}$-0.1 | -0.05691 | -0.4526 to 0.3388 | No | ns | 0.9957 |
| KaiB-1.75, $SasA_{trx}$-0.0 / $SasA_{trx}$-0.3 | -0.2672 | -0.6629 to 0.1285 | No | ns | 0.2498 |
| KaiB-1.75, $SasA_{trx}$-0.0 / $SasA_{trx}$-0.65 | -0.3274 | -0.7231 to 0.06834 | No | ns | 0.1199 |
| KaiB-1.75, $SasA_{trx}$-0.0 / $SasA_{trx}$-1.0 | -0.1683 | -0.5640 to 0.2275 | No | ns | 0.6662 |
| KaiB-1.75, $SasA_{trx}$-0.0 / $SasA_{trx}$-1.8 | -0.08904 | -0.4848 to 0.3067 | No | ns | 0.9653 |
| KaiB-1.75, $SasA_{trx}$-0.0 / $SasA_{trx}$-3.5 | -0.06065 | -0.4564 to 0.3351 | No | ns | 0.9946 |
| KaiA-0.3, $CikA_{psr}$-0.0 / $CikA_{psr}$-0.1 | -0.06478 | -0.3795 to 0.2500 | No | ns | 0.9863 |
| KaiA-0.3, $CikA_{psr}$-0.0 / $CikA_{psr}$-0.3 | -0.06702 | -0.3818 to 0.2477 | No | ns | 0.9833 |
| KaiA-0.3, $CikA_{psr}$-0.0 / $CikA_{psr}$-0.65 | -0.1085 | -0.4233 to 0.2063 | No | ns | 0.851 |
| KaiA-0.3, $CikA_{psr}$-0.0 / $CikA_{psr}$-1.0 | -0.09977 | -0.4145 to 0.2150 | No | ns | 0.892 |
| KaiA-0.3, $CikA_{psr}$-0.0 / $CikA_{psr}$-1.8 | -0.05927 | -0.3740 to 0.2555 | No | ns | 0.9918 |
| KaiA-0.3, $CikA_{psr}$-0.0 / $CikA_{psr}$-3.5 | -0.134 | -0.4487 to 0.1808 | No | ns | 0.7026 |
| KaiA-0.3, $CikA_{psr}$-0.0 / $CikA_{psr}$-5.0 | -0.1477 | -0.4625 to 0.1670 | No | ns | 0.6145 |

Ordinary one-way ANOVA performed with Dunnett's multiple comparisons test was used to compare amplitude values of the FP-PTO under specified conditions: ns, not significant; *, $P < 0.05$; **, $P < 0.01$; ***, $P < 0.001$; #, $P < 0.0001$. If not specified, FP-PTO assays were run with 1.2 μM KaiA, 3.45 μM KaiB, 3.5 μM KaiC, and 0.05 μM fluorescently-labeled KaiB probe.

**Table S6. Ordinary two-way ANOVA of period values from FP-PTO with pairwise comparisons showing how core oscillator components modulate SasA or CikA effects.**

| [Fixed protein-concentration] protein-concentration comparison with / altered protein-concentration (μM) | Predicted (LS) mean diff. | 95.00% CI of diff. | Significant ? | Summary | Adjusted P Value | N1 | N2 |
|---|---|---|---|---|---|---|---|
| [SasA-0.0] KaiB-3.5 / KaiB-1.75 | -1.666 | -2.798 to -0.5337 | Yes | ** | 0.0021 | 7 | 12 |
| [SasA-0.0] KaiB-3.5 / KaiB-7.0 | 0.5423 | -0.8515 to 1.936 | No | ns | 0.6753 | 7 | 5 |
| [SasA-0.0] KaiB-3.5 / KaiB-17.5 | 1.244 | -0.1495 to 2.638 | No | ns | 0.0907 | 7 | 5 |
| [SasA-0.1] KaiB-3.5 / KaiB-1.75 | 0.02 | -1.493 to 1.533 | No | ns | >0.9999 | 5 | 5 |
| [SasA-0.1] KaiB-3.5 / KaiB-7.0 | 0.808 | -0.9387 to 2.555 | No | ns | 0.5642 | 5 | 3 |
| [SasA-0.1] KaiB-3.5 / KaiB-17.5 | 1.228 | -0.5187 to 2.975 | No | ns | 0.2316 | 5 | 3 |
| [SasA-0.3] KaiB-3.5 / KaiB-1.75 | -0.3765 | -1.983 to 1.230 | No | ns | 0.9059 | 5 | 4 |
| [SasA-0.3] KaiB-3.5 / KaiB-7.0 | 0.826 | -0.9225 to 2.575 | No | ns | 0.5504 | 5 | 3 |
| [SasA-0.3] KaiB-3.5 / KaiB-17.5 | 1.069 | -0.6792 to 2.818 | No | ns | 0.3404 | 5 | 3 |
| [SasA-0.65] KaiB-3.5 / KaiB-1.75 | -0.0475 | -1.654 to 1.559 | No | ns | 0.9997 | 5 | 4 |
| [SasA-0.65] KaiB-3.5 / KaiB-7.0 | 0.56 | -1.189 to 2.309 | No | ns | 0.7967 | 5 | 3 |
| [SasA-0.65] KaiB-3.5 / KaiB-17.5 | 1.103 | -0.6452 to 2.852 | No | ns | 0.3152 | 5 | 3 |
| [SasA-1.0] KaiB-3.5 / KaiB-1.75 | -0.174 | -1.687 to 1.339 | No | ns | 0.9868 | 5 | 5 |
| [SasA-1.0] KaiB-3.5 / KaiB-7.0 | 1.041 | -0.7054 to 2.788 | No | ns | 0.3594 | 5 | 3 |
| [SasA-1.0] KaiB-3.5 / KaiB-17.5 | 0.8813 | -0.8654 to 2.628 | No | ns | 0.4959 | 5 | 3 |
| [SasA-1.8] KaiB-3.5 / KaiB-1.75 | 2 | -0.2738 to 4.274 | No | ns | 0.0925 | 2 | 2 |
| [SasA-1.8] KaiB-3.5 / KaiB-7.0 | 3.375 | 0.5902 to 6.160 | Yes | * | 0.0149 | 2 | 1 |
| [CikA-0.0] KaiA-1.2 / KaiA-0.3 | -5.433 | -6.313 to -4.552 | Yes | # | <0.0001 | 7 | 6 |
| [CikA-0.0] KaiA-1.2 / KaiA-0.6 | -1.293 | -2.220 to -0.3661 | Yes | ** | 0.0031 | 7 | 5 |
| [CikA-0.0] KaiA-1.2 / KaiA-2.4 | 0.5891 | -0.3376 to 1.516 | No | ns | 0.3347 | 7 | 5 |
| [CikA-0.0] KaiA-1.2 / KaiA-3.6 | 0.9321 | -0.05985 to 1.924 | No | ns | 0.0719 | 7 | 4 |
| [CikA-0.1] KaiA-1.2 / KaiA-0.3 | -4.394 | -5.728 to -3.060 | Yes | # | <0.0001 | 5 | 2 |
| [CikA-0.1] KaiA-1.2 / KaiA-0.6 | -1.441 | -2.605 to -0.2764 | Yes | ** | 0.0098 | 5 | 3 |
| [CikA-0.1] KaiA-1.2 / KaiA-2.4 | 0.4493 | -0.7150 to 1.614 | No | ns | 0.7689 | 5 | 3 |
| [CikA-0.1] KaiA-1.2 / KaiA-3.6 | -0.174 | -1.920 to 1.572 | No | ns | 0.9979 | 5 | 1 |
| [CikA-0.3] KaiA-1.2 / KaiA-0.3 | -2.871 | -4.205 to -1.537 | Yes | # | <0.0001 | 5 | 2 |
| [CikA-0.3] KaiA-1.2 / KaiA-0.6 | -1.113 | -2.277 to 0.05163 | No | ns | 0.0659 | 5 | 3 |
| [CikA-0.3] KaiA-1.2 / KaiA-2.4 | 0.3707 | -0.7936 to 1.535 | No | ns | 0.8669 | 5 | 3 |
| [CikA-0.3] KaiA-1.2 / KaiA-3.6 | 0.274 | -1.472 to 2.020 | No | ns | 0.9885 | 5 | 1 |
| [CikA-0.65] KaiA-1.2 / KaiA-0.3 | -1.215 | -2.209 to -0.2202 | Yes | * | 0.012 | 5 | 3 |
| [CikA-0.65] KaiA-1.2 / KaiA-0.6 | -0.908 | -1.902 to 0.08644 | No | ns | 0.0824 | 5 | 3 |
| [CikA-0.65] KaiA-1.2 / KaiA-2.4 | 0.222 | -0.7724 to 1.216 | No | ns | 0.9194 | 5 | 3 |
| [CikA-1.0] KaiA-1.2 / KaiA-0.3 | -1.308 | -2.302 to -0.3136 | Yes | ** | 0.0061 | 5 | 3 |
| [CikA-1.0] KaiA-1.2 / KaiA-0.6 | -0.5813 | -1.576 to 0.4131 | No | ns | 0.3817 | 5 | 3 |
| [CikA-1.0] KaiA-1.2 / KaiA-2.4 | 0.05533 | -0.9391 to 1.050 | No | ns | 0.9985 | 5 | 3 |
| [CikA-1.8] KaiA-1.2 / KaiA-0.3 | -0.55 | -1.713 to 0.6129 | No | ns | 0.6158 | 5 | 3 |
| [CikA-1.8] KaiA-1.2 / KaiA-0.6 | -0.87 | -2.033 to 0.2929 | No | ns | 0.2068 | 5 | 3 |

| | | | | | | | |
|---|---|---|---|---|---|---|---|
| [CikA-1.8] KaiA-1.2 / KaiA-2.4 | -0.04667 | -1.210 to 1.116 | No | ns | 0.9999 | 5 | 3 |
| [CikA-1.8] KaiA-1.2 / KaiA-3.6 | -1.51 | -3.254 to 0.2344 | No | ns | 0.1114 | 5 | 1 |
| [CikA-3.5] KaiA-1.2 / KaiA-0.3 | 0.1247 | -0.8698 to 1.119 | No | ns | 0.9839 | 5 | 3 |
| [CikA-3.5] KaiA-1.2 / KaiA-0.6 | -0.1687 | -1.163 to 0.8258 | No | ns | 0.962 | 5 | 3 |
| [CikA-3.5] KaiA-1.2 / KaiA-2.4 | 0.3213 | -0.6731 to 1.316 | No | ns | 0.7969 | 5 | 3 |
| [CikA-5.0] KaiA-1.2 / KaiA-0.3 | 0.1613 | -1.002 to 1.324 | No | ns | 0.9926 | 5 | 3 |
| [CikA-5.0] KaiA-1.2 / KaiA-0.6 | -0.1953 | -1.358 to 0.9676 | No | ns | 0.9847 | 5 | 3 |
| [CikA-5.0] KaiA-1.2 / KaiA-2.4 | -0.132 | -1.295 to 1.031 | No | ns | 0.9965 | 5 | 3 |
| [CikA-5.0] KaiA-1.2 / KaiA-3.6 | -4.302 | -6.046 to -2.558 | Yes | # | <0.0001 | 5 | 1 |

Ordinary two-way ANOVA performed with Dunnett's multiple comparisons test was used to compare pairwise period values of the FP-PTO under specified conditions: ns, not significant; *, $P < 0.05$; **, $P < 0.01$; ***, $P < 0.001$; #, $P < 0.0001$. Predicted least squared (LS) mean difference is reported instead of mean due to comparing means with different number of independent experiments. N1 and N2 specify the number of independent experiments that were able to be fit by FFT-NLLS with the online BioDare suite for the first and second conditions being compared, respectively (6,7). If not specified, oscillators were run with 1.2 μM KaiA, 3.45 μM KaiB, 3.5 μM KaiC, and 0.05 μM fluorescently-labeled KaiB probe.

**Table S7. Ordinary two-way ANOVA of amplitude values from FP-PTO with pairwise comparisons showing how core oscillator components modulate SasA or CikA effects.**

| [Fixed protein-concentration] protein-concentration comparison with / altered protein-concentration (μM) | Predicted (LS) mean diff. | 95.00% CI of diff. | Significant ? | Summary | Adjusted P Value | N1 | N2 |
|---|---|---|---|---|---|---|---|
| [SasA-0.0] KaiB-3.5 / KaiB-1.75 | 0.434 | 0.3145 to 0.5535 | Yes | # | <0.0001 | 7 | 12 |
| [SasA-0.0] KaiB-3.5 / KaiB-7.0 | 0.04656 | -0.1006 to 0.1937 | No | ns | 0.7909 | 7 | 5 |
| [SasA-0.0] KaiB-3.5 / KaiB-17.5 | 0.08743 | -0.05971 to 0.2346 | No | ns | 0.3543 | 7 | 5 |
| [SasA-0.1] KaiB-3.5 / KaiB-1.75 | 0.04169 | -0.1180 to 0.2014 | No | ns | 0.8745 | 5 | 5 |
| [SasA-0.1] KaiB-3.5 / KaiB-7.0 | 0.06957 | -0.1148 to 0.2540 | No | ns | 0.7056 | 5 | 3 |
| [SasA-0.1] KaiB-3.5 / KaiB-17.5 | 0.1301 | -0.05432 to 0.3145 | No | ns | 0.2293 | 5 | 3 |
| [SasA-0.3] KaiB-3.5 / KaiB-1.75 | -0.1321 | -0.3016 to 0.03751 | No | ns | 0.164 | 5 | 4 |
| [SasA-0.3] KaiB-3.5 / KaiB-7.0 | 0.04368 | -0.1409 to 0.2283 | No | ns | 0.9036 | 5 | 3 |
| [SasA-0.3] KaiB-3.5 / KaiB-17.5 | 0.07517 | -0.1094 to 0.2598 | No | ns | 0.6589 | 5 | 3 |
| [SasA-0.65] KaiB-3.5 / KaiB-1.75 | -0.215 | -0.3845 to -0.04541 | Yes | ** | 0.0089 | 5 | 4 |
| [SasA-0.65] KaiB-3.5 / KaiB-7.0 | -0.05956 | -0.2442 to 0.1250 | No | ns | 0.7932 | 5 | 3 |
| [SasA-0.65] KaiB-3.5 / KaiB-17.5 | 0.01352 | -0.1711 to 0.1981 | No | ns | 0.9966 | 5 | 3 |
| [SasA-1.0] KaiB-3.5 / KaiB-1.75 | -0.1468 | -0.3065 to 0.01293 | No | ns | 0.0792 | 5 | 5 |
| [SasA-1.0] KaiB-3.5 / KaiB-7.0 | 0.04559 | -0.1388 to 0.2300 | No | ns | 0.8907 | 5 | 3 |
| [SasA-1.0] KaiB-3.5 / KaiB-17.5 | 0.03213 | -0.1523 to 0.2165 | No | ns | 0.9571 | 5 | 3 |
| [SasA-1.8] KaiB-3.5 / KaiB-1.75 | 0.01854 | -0.2246 to 0.2617 | No | ns | 0.9792 | 2 | 2 |
| [SasA-1.8] KaiB-3.5 / KaiB-7.0 | -0.01082 | -0.3086 to 0.2869 | No | ns | 0.9952 | 2 | 1 |
| [CikA-0.0] KaiA-1.2 / KaiA-0.3 | 0.3983 | 0.2520 to 0.5447 | Yes | # | <0.0001 | 7 | 5 |
| [CikA-0.0] KaiA-1.2 / KaiA-0.6 | 0.1897 | 0.04336 to 0.3361 | Yes | ** | 0.0066 | 7 | 5 |
| [CikA-0.0] KaiA-1.2 / KaiA-2.4 | 0.12 | -0.02640 to 0.2663 | No | ns | 0.1407 | 7 | 5 |
| [CikA-0.0] KaiA-1.2 / KaiA-3.6 | 0.2376 | 0.08098 to 0.3943 | Yes | ** | 0.0012 | 7 | 4 |
| [CikA-0.1] KaiA-1.2 / KaiA-0.3 | 0.3469 | 0.1364 to 0.5574 | Yes | *** | 0.0004 | 5 | 2 |
| [CikA-0.1] KaiA-1.2 / KaiA-0.6 | 0.1197 | -0.06407 to 0.3034 | No | ns | 0.325 | 5 | 3 |
| [CikA-0.1] KaiA-1.2 / KaiA-2.4 | 0.0546 | -0.1291 to 0.2383 | No | ns | 0.8923 | 5 | 3 |
| [CikA-0.1] KaiA-1.2 / KaiA-3.6 | 0.2974 | 0.02178 to 0.5730 | Yes | * | 0.0299 | 5 | 1 |
| [CikA-0.3] KaiA-1.2 / KaiA-0.3 | 0.3029 | 0.09246 to 0.5134 | Yes | ** | 0.0021 | 5 | 2 |
| [CikA-0.3] KaiA-1.2 / KaiA-0.6 | 0.08615 | -0.09757 to 0.2699 | No | ns | 0.626 | 5 | 3 |
| [CikA-0.3] KaiA-1.2 / KaiA-2.4 | 0.06877 | -0.1150 to 0.2525 | No | ns | 0.7871 | 5 | 3 |
| [CikA-0.3] KaiA-1.2 / KaiA-3.6 | 0.4703 | 0.1947 to 0.7459 | Yes | *** | 0.0002 | 5 | 1 |
| [CikA-0.65] KaiA-1.2 / KaiA-0.3 | 0.3491 | 0.1872 to 0.5111 | Yes | # | <0.0001 | 5 | 3 |
| [CikA-0.65] KaiA-1.2 / KaiA-0.6 | 0.06287 | -0.09909 to 0.2248 | No | ns | 0.6955 | 5 | 3 |
| [CikA-0.65] KaiA-1.2 / KaiA-2.4 | 0.08232 | -0.07964 to 0.2443 | No | ns | 0.4976 | 5 | 3 |
| [CikA-1.0] KaiA-1.2 / KaiA-0.3 | 0.2408 | 0.07883 to 0.4027 | Yes | ** | 0.0017 | 5 | 3 |
| [CikA-1.0] KaiA-1.2 / KaiA-0.6 | 0.06642 | -0.09553 to 0.2284 | No | ns | 0.6594 | 5 | 3 |
| [CikA-1.0] KaiA-1.2 / KaiA-2.4 | 0.0411 | -0.1209 to 0.2031 | No | ns | 0.8874 | 5 | 3 |
| [CikA-1.8] KaiA-1.2 / KaiA-0.3 | 0.1746 | -0.008936 to 0.3581 | No | ns | 0.0675 | 5 | 3 |
| [CikA-1.8] KaiA-1.2 / KaiA-0.6 | -0.03042 | -0.2139 to 0.1531 | No | ns | 0.9854 | 5 | 3 |

| | | | | | | | |
|---|---|---|---|---|---|---|---|
| [CikA-1.8] KaiA-1.2 / KaiA-2.4 | 0.08793 | -0.09558 to 0.2714 | No | ns | 0.6045 | 5 | 3 |
| [CikA-1.8] KaiA-1.2 / KaiA-3.6 | 0.262 | -0.01327 to 0.5373 | No | ns | 0.0673 | 5 | 1 |
| [CikA-3.5] KaiA-1.2 / KaiA-0.3 | 0.1377 | -0.02421 to 0.2997 | No | ns | 0.1154 | 5 | 3 |
| [CikA-3.5] KaiA-1.2 / KaiA-0.6 | -0.03605 | -0.1980 to 0.1259 | No | ns | 0.92 | 5 | 3 |
| [CikA-3.5] KaiA-1.2 / KaiA-2.4 | 0.1386 | -0.02338 to 0.3005 | No | ns | 0.1124 | 5 | 3 |
| [CikA-5.0] KaiA-1.2 / KaiA-0.3 | 0.09614 | -0.08737 to 0.2796 | No | ns | 0.5253 | 5 | 3 |
| [CikA-5.0] KaiA-1.2 / KaiA-0.6 | -0.02335 | -0.2069 to 0.1602 | No | ns | 0.9947 | 5 | 3 |
| [CikA-5.0] KaiA-1.2 / KaiA-2.4 | 0.07869 | -0.1048 to 0.2622 | No | ns | 0.6934 | 5 | 3 |
| [CikA-5.0] KaiA-1.2 / KaiA-3.6 | 0.2078 | -0.06747 to 0.4830 | No | ns | 0.1999 | 5 | 1 |

Ordinary two-way ANOVA performed with Dunnett's multiple comparisons test was used to compare pairwise amplitude values of the FP-PTO under specified conditions: ns, not significant; *, $P < 0.05$; **, $P < 0.01$; ***, $P < 0.001$; #, $P < 0.0001$. Predicted least squared (LS) mean difference is reported instead of mean due to comparing means with different number of independent experiments. N1 and N2 specify the number of independent experiments that were able to be fit by FFT-NLLS with the online BioDare suite for the first and second conditions being compared, respectively (6,7). If not specified, oscillators were run with 1.2 μM KaiA, 3.45 μM KaiB, 3.5 μM KaiC, and 0.05 μM fluorescently-labeled KaiB probe.

### Table S8. Plasmids and primers used in generating cyanobacterial strains.

| Plasmids | Description | Source |
|---|---|---|
| pSL2680 | CRISPR/Cas12a plasmid; Km resistance | Addgene (#85581) |
| pSL2680-HA | pSL2680 + SasA H28A substitution | This study |
| pSL2680-NA | pSL2680 + SasA N93A substitution | This study |
| pSL2680-NE | pSL2680 + SasA N93E substitution | This study |
| pSL2680-94 | pSL2680 + SasA Q94A substitution | This study |
| pSL2680-HA/94 | pSL2680 + SasA H28A and Q94A substitutions | This study |
| pSL2680-97 | pSL2680 + SasA Q97A substitution | This study |
| pSL2680-QE | pSL2680 + SasA Q97E substitution | This study |
| pSL2680-101 | pSL2680 + SasA Q101A substitution | This study |

| Primers | Sequence (5'-3') |
|---|---|
| **Primers used for pSL2680-HA** | |
| H28A gRNA F | AGATTGCAGCGGGTTAAAAATATT |
| H28A gRNA R | AGACAATATTTTTAACCCGCTGCA |
| H28A homology arm upstream F | TAGCTTTAATGCGGTAGTTGGTACCATGATCGACGCCTGTCGA |
| H28A homology arm upstream R | TTTAACCCGCTGCACGATGGCCTGTGACAGGGGCCG |
| H28A homology arm downstream F | CGGCCCCTGTCACAGGCCATCGTGCAGCGGGTTAAA |
| **Primers used for pSL2680-NA** | |
| N93A gRNA F | AGATGCTAATTGATCGGTGAGGTC |
| N93A gRNA R | AGACGACCTCACCGATCAATTAGC |
| N93A homology arm upstream F | CATTTTTTTGTCTAGCTTTAATGCGGTAGTTGGTACC CTGGCGATGGACTTGCACTCA |
| N93A homology arm upstream R | CTGGGGCAACTGGGCGGCTAATTGATCGGT |
| N93A homology arm downstream F | ACCGATCAATTAGCCGCCCAGTTGCCCCAG |
| N93A homology arm downstream R | GCCCGGATTACAGATCCTCTAGAGTCGACGGTACC TTAGCAGGGCATGGTGTAGC |
| **Primers used for pSL2680-NE** | |
| N93E gRNA F | AGATGCTAATTGATCGGTGAGGTC |
| N93E gRNA R | AGACGACCTCACCGATCAATTAGC |
| N93E homology arm upstream F | CATTTTTTTGTCTAGCTTTAATGCGGTAGTTGGTACC CTGGCGATGGACTTGCACTCA |
| N93E homology arm upstream R | CTGGGGCAACTGCTCGGCTAATTGATCGGT |
| N93E homology arm downstream F | ACCGATCAATTAGCCGAGCAGTTGCCCCAG |
| N93E homology arm downstream R | GCCCGGATTACAGATCCTCTAGAGTCGACGGTACC TTAGCAGGGCATGGTGTAGC |

| | |
|---|---|
| **Primers used for pSL2680-94** | |
| Q94A gRNA F | AGATGTTGGCTAATTGATCGGTGA |
| Q94AgRNA R | AGACTCACCGATCAATTAGCCAAC |
| Q94A homology arm upstream F | CATTTTTTTGTCTAGCTTTAATGCGGTAGTTGGTACC CTGGCGATGGACTTGCACTCA |
| Q94A homology arm upstream R | CCACTGGGGCAACGCGTTGGCTAATTG |
| Q94A homology arm downstream F | CAATTAGCCAACGCGTTGCCCCAGTGG |
| Q94A homology arm downstream R | GCCCGGATTACAGATCCTCTAGAGTCGACGGTACC TTAGCAGGGCATGGTGTAGC |
| **Primers used for pSL2680 HA/94** | |
| HA/94 gRNA F | AGATACGAAGAAAGCTCAGTGAGC |
| HA/94 gRNA R | AGACGCTCACTGAGCTTTCTTCGT |
| HA/94 homology arm upstream F | CATTTTTTTGTCTAGCTTTAATGCGGTAGTTGGTACC CTGGCGATGGACTTGCACTCA |
| HA/94 homology arm upstream R | GCTCACTGAGCTTTCTTCGTGTATCCGCCAAATTGT |
| HA94 homology arm downstream F | ACAATTTGGCGGATACACGAAGAAAGCTCAGTGAGC |
| HA94 homology arm downstream R | CAGATCCTCTAGAGTCGACGGTACC ATCGTGCCTGATCGAACA |
| **Primers used for pSL2680-97** | |
| Q97A gRNA F | AGATGGGCAACTGGTTGGCTAATT |
| Q97A gRNA R | AGACAATTAGCCAACCAGTTGCCC |
| Q97A homology arm upstream F | CATTTTTTTGTCTAGCTTTAATGCGGTAGTTGGTACC CTGGCGATGGACTTGCACTCA |
| Q97A homology arm upstream R | CTGAACCAGCCACGCGGGCAACTGGTT |
| Q97A homology arm downstream F | AACCAGTTGCCCGCGTGGCTGGTTCAG |
| Q97A homology arm downstream R | GCCCGGATTACAGATCCTCTAGAGTCGACGGTACC TTAGCAGGGCATGGTGTAGC |
| **Primers used for pSL2680-QE** | |
| Q97E gRNA F | AGATTGAGTGGCATCGACCTCACC |
| Q97E gRNA R | AGACGGTGAGGTCGATGCCACTCA |
| Q97E homology arm upstream F | CATTTTTTTGTCTAGCTTTAATGCGGTAGTTGGTACC CTGGCGATGGACTTGCACTCA |
| Q97E homology arm upstream R | GGCTAATTGATCGGTGAGGTCGATGCCACTCAGCACTTGGC |
| Q97E homology arm downstream F | ACCTCACCGATCAATTAGCCAACCAGTTGCCCGAGTGGCTGG |
| Q97E homology arm downstream R | GCCCGGATTACAGATCCTCTAGAGTCGACGGTACC TTAGCAGGGCATGGTGTAGC |

| | |
|---|---|
| **Primers used for pSL2680-101** | |
| Q101A gRNA F | AGATAACCAGCCACTGGGGCAACT |
| Q101A gRNA R | AGACAGTTGCCCCAGTGGCTGGTT |
| Q101A homology arm upstream F | CATTTTTTTGTCTAGCTTTAATGCGGTAGTTGGTACC CTGGCGATGGACTTGCACTCA |
| Q101A homology arm upstream R | AAAGGCCTCTTGCGCAACCAGCCACTG |
| Q101A homology arm downstream F | CAGTGGCTGGTTGCGCAAGAGGCCTTT |
| Q101A homology arm downstream R | GCCCGGATTACAGATCCTCTAGAGTCGACGGTACC TTAGCAGGGCATGGTGTAGC |
| **Primers used for colony PCR** | |
| *sasA1_SNP_chkF* | CGAGTTAATGGGAGAGTCTCTGTC |
| *sasA1_SNP_chkR* | GGCCTAGCTCCGTGAACG |

**Table S9. Cyanobacterial strains used in this study.**

| Strain | Genotype (NS denotes neutral site) | Antibiotic resistance | Source |
|---|---|---|---|
| WT (AMC541) | NSII-$P_{kaiBC}$::*luc* | Cm | Lab collection |
| Δ*sasA* (AMC1192) | NSII-$P_{kaiBC}$::*luc* Δ*sasA* | Cm | Lab collection |
| *sasA*-H28A | NSII-$P_{kaiBC}$::*luc*. *sasA*-H28A | Cm | This study |
| *sasA*-N93A | NSII-$P_{kaiBC}$::*luc*. *sasA*-N93A | Cm | This study |
| *sasA*-N93E | NSII-$P_{kaiBC}$::*luc*. *sasA*-N93E | Cm | This study |
| *sasA*-Q94A | NSII-$P_{kaiBC}$::*luc* *sasA*-Q94A | Cm | This study |
| *sasA*-H28A/Q94A | NSII-$P_{kaiBC}$::*luc* *sasA*-H28A/Q94A | Cm | This study |
| *sasA*-Q97A | NSII-$P_{kaiBC}$::*luc*. *sasA*-Q97A | Cm | This study |
| *sasA*-Q97E | NSII-$P_{kaiBC}$::*luc*. *sasA*-Q97E | Cm | This study |
| *sasA*-Q101A | NSII-$P_{kaiBC}$::*luc* *sasA*-Q101A | Cm | This study |

**Data S1. DynaFit script for modeling of 2D titration assays (separate file)**
Text file containing annotated DynaFit script for two-site binding model.

**Data S2. DynaFit script for simulation of 2D titration assays (separate file)**
Text file containing annotated DynaFit scripts to analyze 2D titration assays.

**Data S3. Input/output data from least-squares thermodynamic modeling (separate file)**
Raw and analyzed datasets used for thermodynamic modeling.

**Data S4. FP-PTO raw data file (separate file)**
Data organized by figures of interest.

**Data S5. FP-PTO normalized data file (separate file)**
Data organized by figures of interest.

**Data S6. BioDare2 quantitative analysis of oscillatory rhythm under various conditions data file (separate file)**
Complete analysis of all FP-PTO assay rhythms containing period, amplitude, and phase determinations.
