## Supplementary material for "Structural mimicry confers robustness in the cyanobacterial circadian clock": Data S1

DynaFit (BioKin) script for global fitting of KaiC binding to fluorescent KaiB in the presence of an additive.

http://www.biokin.com/dynafit/.

Assay monitored using fluorescence anisotropy

B = KaiB probe

C = KaiC (enter as dimer concentration)

S = additive such as SasA

Input data must be formatted as a .csv file with no headers.

Usually it is best to constrain the fit initially using a miniumum number of parameters to fit the general profile of the titration,
and then float the other parameters in turn and check the fit still converges.

To float a parameter, append “?” after the initial value.

Once the program converges to a satisfactory solution, 
remove the ; (comment) in front of "confidence = monte-carlo" and re-run
to assess any covariance of the parameters and determne the 95% confidence interavals.

; _______________________________________________________
[task]
   task = fit
   data = equilibria
   ;confidence = monte-carlo  ; uncomment this to run monte-carlo analysis
 
[mechanism] 
   B + C <==> BC   	:     K1     dissoc
   B + BC  <==> B2C   	:     K5     dissoc

  
   S + C <==> SC   	:     K2     dissoc
   S + SC  <==> S2C     	:     K6     dissoc
 
   S + BC   <==> SBC    	:     K4      dissoc

  ;B + SC <==> SBC           :     K3      dissoc   keep this as a comment statement as K3 defined by other constants

 
[constants]
   K1 = 0.466? 	;initial estimate determined from fit to data in absence of additive (SasA)
   K5  = 4 * K1  	;macroscopic constant for dimer with independent sites Float this parameter to assess homotropic cooperativity
  
   K2 = 0.01 ?
   K6 = 4 * K2   	;macroscopic constant for dimer with independent sites Float this parameter to assess homotropic cooperativity
 
   K4  = .0003?

  ; K3 = K1*K4/K2  for thermodynalic balance

  	
[concentrations]
   B = 0.05

[responses]
   intensive ; fluorescence anisotropy - float these and then define prior running monte-carlo
	     
   B = 0.140
   BC = 0.185
   B2C = 0.211
  
   SBC = 0.211 ; average fit from full 2D titrations was 
	       ; 0.211 for SasA
	       ; 0.203 for fsKaiB 

  
[data]
   variable  C
   plot logarithmic

   directory ./DynaFit_scripts/
   sheet     datafilename.csv 
	; Enter KaiC concentration as dimer in column 1 and anisotropy subsequent columns. All columns called must be populated.
	; To run with datasets with no or fewer additive concentrations, comment out some of the below 

   columns 2 | conc B = 0.05   |conc S = 0 	| label SasA=0
   columns 4 | conc B = 0.05  |conc S = 0.05 	| label SasA=0.05
   columns 6 | conc B = 0.05  |conc S = 0.1 	| label SasA=0.1
   columns 8 | conc B = 0.05  |conc S = 0.3 	| label SasA=0.3
   columns 10 | conc B = 0.05  |conc S = 0.6 	| label SasA=0.6
   columns 12 | conc B = 0.05  |conc S = 1.0	| label SasA=1 
   columns 14 | conc B = 0.05  |conc S = 3.5	| label SasA=3.5 
   
   
; conc S values above 0.3 may be floated to allow for experimental errors in relative concentrations of SasA and KaiC

[output]
   directory ./DynaFit_scripts/

[end]
