## Supplementary material for "Structural mimicry confers robustness in the cyanobacterial circadian clock": Data S2

DynaFit (BioKin) script for simulation of KaiC binding to fluorescent KaiB in the presence of an additive.

http://www.biokin.com/dynafit/.

See additional comments in script for fitting data

DynaFit modifies the procedure for simulation but still requires a dummy data set to define concentrations of additive S

Same data formatting rules apply as in 
;______________________________________________________________________
[task]
   task = simulate
   data = equilibria
  ; confidence = monte-carlo
 
[mechanism] 
   B + C <==> BC   	:     K1     dissoc
   B + BC  <==> B2C   	:     K5     dissoc

  
   S + C <==> SC   	:     K2     dissoc
   S + SC  <==> S2C     	:     K6      dissoc
 
   S + BC   <==> SBC    	:     K4      dissoc

 
[constants]
   K1 = 1 ; from rounded fit to data in absence of SasA
   K5  = 4 * K1  ; macroscopic constant for dimer
  
   K2 = 0.002 
   K6  = 4 * K2   ; non-homo-cooperative binding
 
   K4  = 0.00002   ; heterotropic cooperativity index = K1/K3 = K2/K4
		    ; alter K4 to adjust cooperativity index (c.i.) in the simulation
		    ; 0.002 for "c.i. = 1"
		    ; 0.0002 for "c.i. = 10"
		    ; 0.00002 for "c.i. = 100" 

  	
[concentrations]
   B = 0.05
[responses]
   intensive ; fluorescence anisotropy values based of fitted data
   B = 0.14 
   BC = 0.18 
   B2C = 0.18
  
   SBC = 0.21

  
[data]
   variable  C
   mesh logarithmic from 0.001 to 1000 step 2     ; additional statement for simulation to define concentration range of KaiC
   plot logarithmic

   directory ./DynaFit_scripts/
   sheet     data_filename.csv ;used to define S range in simulation
   
columns 3 | conc B = 0.05   |conc S = 0 	| label SasA=0
   columns 4 | conc B = 0.05  |conc S = 0.05 	| label SasA=0.05
   columns 5 | conc B = 0.05  |conc S = 0.1 	| label SasA=0.1
   columns 6 | conc B = 0.05  |conc S = 0.3 	| label SasA=0.3
   columns 7 | conc B = 0.05  |conc S = 0.6 	| label SasA=0.6
   columns 8 | conc B = 0.05  |conc S = 1	| label SasA=1
   columns 11 | conc B = 0.05  |conc S = 3.5	| label SasA=3.5
   
[output]
   directory ./DynaFit_scripts/


[end]
